# Optimising Elastic Network Models for Protein Dynamics and Allostery: Spatial and Modal Cut-offs and Backbone Stiffness

**DOI:** 10.1101/2022.05.13.491757

**Authors:** Igors Dubanevics, Tom C.B. McLeish

## Abstract

The family of coarse-grained models for protein dynamics known as Elastic Network Models (ENMs) require a careful choice of parameters to represent well experimental measurements or fully-atomistic simulations. The most basic ENM that represents each protein residue by a node at the position of its C-alpha atom, all connected by springs of equal stiffness, up to a cut-off in distance. Even at this level, a choice is required of the optimum cut-off distance and the upper limit of elastic normal modes taken in any sum for physical properties, such as dynamic correlation or allosteric effects on binding. Additionally, backbone-enhanced ENM (BENM) may improve the model by allocating a higher stiffness to springs that connect along with the protein backbone. This work reports on the effect of varying these three parameters (distance and mode cutoffs, backbone stiffness) on the dynamical structure of three proteins, Catabolite Activator Protein (CAP), Glutathione S-transferase (GST), and the SARS-CoV- 2 Main Protease (M^pro^). Our main results are: (1) balancing B-factor and dispersion-relation predictions, a near-universal optimal value of 8.5 angstroms is advisable for ENMs; (2) inhomogeneity in elasticity brings the first mode containing spatial structure not well-resolved by the ENM typically within the first 20; (3) the BENM only affects modes in the upper third of the distribution, and, additionally to the ENM, is only able to model the dispersion curve better in this vicinity; (4) BENM does not typically affect fluctuation-allostery, which also requires careful treatment of the effector binding to the host protein to capture.

## Introduction

### Fluctuation-Allostery

Allosteric mechanisms for distant control of binding and activation fall into two main classes: those which invoke significant conformational change (the original scenario of Monod, Wyman and Changeaux [1], and mechanisms that invoke the modification of thermal (entropic) fluctuations about a fixed, mean conformation [2–5]. Such ’fluctuation allostery’ has also been termed ’allostery without conformational change’ but this appellation requires careful qualification, for the lack of conformational change applies at the level of alpha-carbon co-ordinates only. In any case, the mechanism recruits mostly global, low-frequency modes of internal protein motion, which are well-captured by correspondingly coarsegrained mechanical representations of the protein [6, 7]. As an alternative to such a continuum description, the same underlying physics of correlated entropic changes on binding effector and substrate has been described in terms of changes to the (thermal) occupation distribution of multiple states [8]. In all these approaches, allosteric signalling is brought about by the modifications that the effector-binding makes to the dynamic structure of the protein, such that the entropic part of the free-energy of substrate binding is modified in turn. One effective tool that has been successfully used to identify the relevant dynamic structure for fluctuation allostery is the Elastic Network Model (ENM) [9, 10]. The ENM resolves protein structure at the level of alpha-carbon sites only, which are represented as nodes connected by harmonic springs within a fixed cut-off radius from each other. In its original and most basic form, all harmonic bonds are accorded the same stiffness. Local point mutations can be modelled by changing the moduli of springs attached to the corresponding residue, and effector-binding by the addition of nodes (and local harmonic potentials) at the corresponding co-ordinates. The most significant contributions to both the correlated dynamics of distant residues, and to the entropy arising from structural fluctuation, come from global modes, because it is these that meet the requirement of correlated local strain at the (typically distant) effector and allosteric binding sites [4]. The ENM approximation allows straightforward calculation of the dynamics of these low-frequency global modes, since they naturally average out the local variation of bond strengths and densities.We note that this low-frequency mode faithfulness of ENM models is true, even for global modes comprise motions such as relative rotations of entire sub-domains. Although the ENM does not capture the potential for bond rotation that resides at the nexus of such relative motion, the majority of the potential energy for such global modes arises, not from single bonds, but from the sum of many residue-residue distance perturbation at the interface [11]. This observation motivates the weighting of lower modes for the parameterisation of the ENMs in this study, as below. Once ENMs are parameterised so that they agree at low frequencies with molecular dynamics simulations, they can be used to predict other quantities.Such an approach was successfully used to identify candidate control residues whose mutation may control allostery of effector binding in the homodimer transcription factor CAP [12].

The latter study, and others, have shown that, while the huge reduction in the number of degrees of freedom that the ENM constitutes captures neither the quantitative values of free energies, nor the numerical changes to those values on mutation that are seen in experiment, it is able to rank them qualitatively. It is also able to capture the functional form of the thermal dynamics of a protein for a significant set of low-frequency modes, and identify residues whose mutation has a strong effect on the allostery. Naturally, those aspects of protein function that rely on side-chain structures and interactions, or on very fast processes, are not captured by any coarse-grained model, including those of the ENM class. However, low-frequency dynamics, and those processes that modify them, can be addressed by ENM models. In particular, the coarse-graining of the ENM can determine which residues qualify as candidates for allosteric control through mutation. This method has been embedded in, among other available tools, the open-source software (’DDPT’), used in the previous study on allosteric homodimers, and confirmed by experimental calorimetry on model-designed mutations [13].

In terms of applications, the ENM approach to allostery allows, for example, the rapid assessment of the druggability of proteins. This rapid screening is even more significant in the case of constantly evolving diseases, like SARS-CoV-2 (SCoV2). For example, our previous work on the SCoV2 main protease identified active residues that were also experimentally verified, and suggested sites for potential non-competitive inhibition [14]. Another recent study analysed allosteric pathways of the spike protein, a structurally essential and highly mutable part of the viral capsid, and similarly provided suggestions for allosteric drug design [15]. Other work with ENMs has also touched upon mutability, dynamics and evolution of the protein. Increasingly, protein-dynamics research is exploring the interplay between protein dynamics and evolution, with help of the ENM, which allows rapid assessment of protein dynamics and is able to relate it to evolution at the level of individual residues [16, 17]. The essential global-mode structure turns out, for example, to be much more highly conserved through evolution, than high-frequency structure [18].

Even at this level of coarse-graining, which dispenses with the huge space of possible parameterisation of fully-atomistic models, there remain open questions in regard to the optimal parameterisation, the range of physical applicability of ENM models, and possible refinements of the ENM model itself. These have been the subjects of recent reviews and analyses of the ENM approach [19–21]. Potential extensions and suggested refinements include the breaking of some bonds during structural fluctuation [22], the relative importance of refining protein geometry over residue specificity [23], and the introduction of specific residue-residue bond strengths [24]. However, there are even more basic parameter choices routinely made within basic ENM models, such as appropriate cut-offs to the mode sum and in spatial bond-distance, that still lack understanding in terms of their optimal choices. Of especial interest to research in the application areas of protein binding, druggability etc. are calculations of fluctuation-generated allosteric response. In these areas it is important to identify if a consistent, even automated, implementation of coarse-grained models, can reliably be constructed for high-throughput computational screening [25]. In the service of such a goal, this paper reports on further progress in the search for appropriate biological and physical constraints that inform an optimal selection of an ENM for a protein to study its allosteric structure.

### Questions

As noted above, the parameterisation of the traditional versions of ENMs that have been developed in the study of coarse-grained dynamical structure of proteins has typically called on two quantitative choices even in the simplest of models, in which all harmonic springs between the C_*α*_ nodes are identical. The first choice corresponds to the distance within which the ENM constructs a harmonic bond between two residues - an effective distance cut-off for inter-residue interaction. The second recognises that physical and thermodynamic quantities are all expressed as a sum over the dynamic normal modes of the linear elastic system that results from the ENM, but that not all modes are physically-relevant. This is because at some point in the mode-spectrum, eigenmodes begin to contain spatial structures of a size too small to be captured accurately (or spatially resolved) at the ENM level of coarse-graining. Such an effect therefore gives rise to a physically-generated mode cut-off. A third question arises in a slightly more sophisticated versions of the ENM [26] in which backbone connections are accorded a higher strength than all other bonds. This ’backbone-enhanced ENM’ (BENM) requires one additional parameter, the ratio of the two values of spring constant then employed.The last, a fourth question, explores the role of EN ligand modelling in numerical cooperativity computation. Protein-ligand interaction has always been treated separately from protein-protein interaction in MD simulations and requires a careful attention from a simulation carrier, as in case of general AMBER force field (GAFF) [27]. Two types of binding effector (ligand) EN representations are presented here: single-bead and all heavy-atom representations with custom protein-ligand bonding based on the crystallographic data. Thus, the four key parameterisation questions are therefore:

- What distance cutoff to choose?
- What mode cutoff to use?
- How does backbone-enhanced ENM (BENM) affect the protein dynamical structure and numerical calculation of cooperativity?
- What effect does ligand EN representation and its connectivity with protein EN nodes has on numerical cooperativity?

Each of these questions implicates a quantitative parameter. The distance cutoff we will represent by *d*_*c*_ (measured in Å). The mode cutoff in sums for physical quantities will be *n*_*c*_. The backbone stiffness is represented by *ϵ*, the dimensionless ratio of the strengths of backbone springs to others. The set *d*_*c*_, *n*_*c*_, *ϵ* constitutes the parameter space that this work will explore. The answers to the three questions may, of course, depend on the particular protein under study. It is also desirable that the optimising parameters be as near-universal as possible, but given the strong heterogeneity of proteins themselves, as well as the structure differences between them, it is not guaranteed that a single universal optimal set exists. In the case that it does not, a new goal would be to identify, if possible, prior characteristics of proteins that may suggest one or other class of optimum parameters.

Such a programme also requires success-measures for the models that may be constructed by exploring the parameter space, and used to evaluate optima within the *d*_*c*_, *n*_*c*_, *ϵ* parameter space. In the following we will use: (i) the agreement for B-factor values measured in both experiment and by molecular dynamics; (ii) the dispersion relation for normal mode frequencies, in terms of mode number, *ω*(*n*), again comparing against molecular dynamics simulations; (iii) the allosteric free energy or equivalently the ratio of binding constants for holo1 and holo2 forms of the proteins.

We will perform this analysis for three protein homodimers, the first two of which have been subject to ENM analysis before. The proteins of choice are: the Catabolite Activator Protein (CAP) [28], Glutathione S-transferase (GST) [29], and the SARS-CoV-2 Main Protease (M^pro^) [14]. The next section reviews the structure of these proteins and their ligands, and the construction of EMNs for them. There follows a discussion of the results of varying the three key parameters on their dynamic and allosteric structures, before drawing conclusions for the development and the careful future use of ENMs.

### Proteins

This section presents the three protein homodimers chosen as exemplars for optimising ENMs for dynamic and allosteric predictions. They are the chosen subjects for the intensive calculations and analyses devoted to addressing the three main questions identified in the introduction. Structures for the three proteins, CAP, GST, and M^pro^, are given from two perspectives and in their ligand-bound forms in figure 1, as well as showing the detailed structures of the ligands in each case.

**Figure 1:**
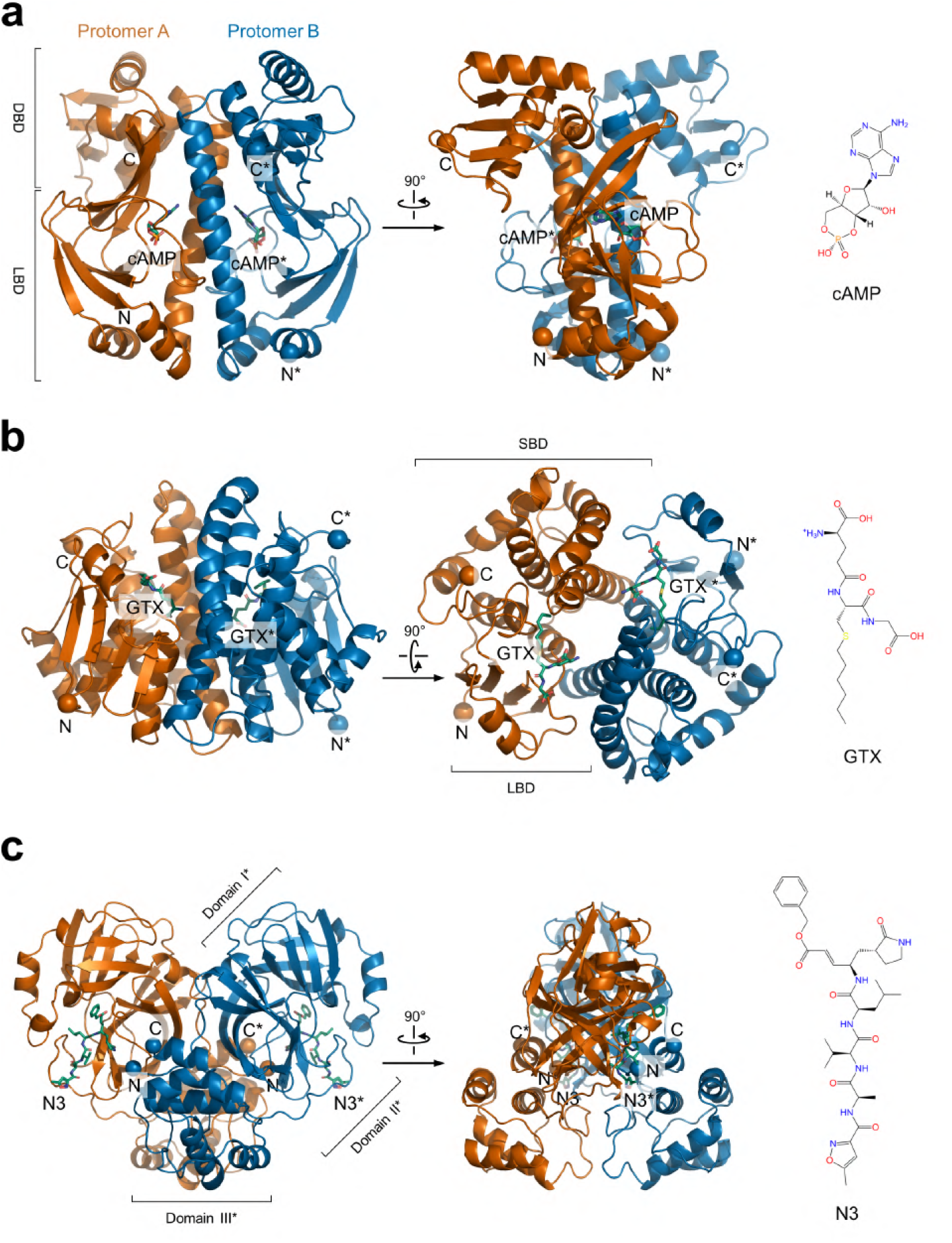
Protein-ligand complexes used in this study. **a**, CAP with bound cAMP (PDB ID: 4HZF). DNA binding domain (DBD) and ligand binding domain (LBD) are shown only for protomer A. **b**, GST in complex with GTX ligand (PDB ID: 1M9A). Substrate binding domain (SBD) and LBD are shown only for protomer A. **c**, SCoV2 M^pro^ with N3 inhibitor (PDB ID: 7BQY). Three protein domains are labelled only for protomer B. Protomer A is in *blue* while protomer B in *orange*. Asterisks mark residues from protomer B.

Table 1 gives detailed information on the PDB files, sizes and molecular weights of the target proteins. In the following, we give more detailed discussion of the structure, function and biological context of each protein, and the motivation for its choice in this study.

**Table 1:** Protein-ligand complexes used in this study. Protein section: Protein Data Bank accession ID; abbreviation; protomer IDs and residue numbers; number of alpha-carbons; trimmed residue structure for ENM’ number of elastic network beads. Ligand section: abbreviation, number of heavy atoms (non-hydrogen); total atomic mass of a ligand.

| PDB ID | Protein |  |  |  |  | Ligand |  |  |
| --- | --- | --- | --- | --- | --- | --- | --- | --- |
| | Name | Protomers | $C_\alpha$ (#) | ENM protomers | EN nodes (#) | Name | Atoms (#) | Mass (amu) |
| 4HZF | CAP | A: 9-208 | 400 | A: 11-207 | 394 | cAMP | 34 | 329.21 |
|  |  | B: 10-209 |  | B: 11-207 |  |  |  |  |
| 1M9A | GST | A: 1-216 | 432 | A: 4-213 | 420 | GTX | 56 | 392.49 |
|  |  | B: 1-216 |  | B: 4-213 |  |  |  |  |
| 7BQY | M<sup>Pro</sup> | A: 1-301 | 602 | A: 2-300 | 598 | N3 | 97 | 680.79 |
|  |  | B: 1-301 |  | B: 2-300 |  |  |  |  |

### Catabolite Activator Protein

The Catabolite Activator Protein (CAP), often referred to as the cAMP receptor protein, is an extensively studied transcription factor native to certain strains of *E. coli* [30]. It is known to bind to a promoter of the *lac* operon gene as well up to 100 other promoters.

In solution CAP exists as a homodimer with two-fold *C*_2_ symmetry; each subunit consisting of 210 amino acids (Figure 1). The protomer contains two domains: the N-terminal 138 amino acids constitutes the ligand-binding domain (LBD); the C-terminal portion of the molecule the DNA binding domain (DBD). In its apo form (without the ligand cAMP bound) it is inactive, but when two molecules of cAMP bind to the dimer they activate the protein and increases its affinity for binding to DNA sufficient to ensure predominance of the bound state in the living organism.

Experimental work from more than 15 years ago first showed dynamically driven allostery in CAP [31], followed by several studies which verified the finding [12, 32–35]. Negative allostery between the CAP’s cAMP binding sites is of the order of *K*_2_/*K*_1_ ≈ 1.7 (*K*_1_ and *K*_2_ represent the dissociation constants for the first and second ligand binding events).

Through studies of apo, single ligand bound (holo1), and double ligand bound (holo2) CAP states by NMR and isothermal calorimetry (ITC) it was demonstrated that there were minimal changes in protein structure between the single and double-liganded proteins. Conversely, there were distinct changes in protein motions between the three binding states, and the enhanced negative cooperativity was demonstrated to be driven by the associated changes in entropy. Binding of the first ligand molecule had little effect on motion in the ps–ns range but activated slow motions (µs–ms regime) across both protomers [31]. Binding of the second ligand molecule suppressed both fast and slow motions. The suppression of the µs–ms range motions on binding the second ligand molecule represents the entropic penalty that is the source of negative co-operativity. This study demonstrated that allostery can arise through a purely dynamic mechanism. Further investigation used ENM models to identify residues whose mutations could most sensitively affect CAP’s allosteric properties, which were qualitatively confirmed experimentally [12].

### Glutathione S-transferase

Glutathione transferases (GSTs) constitute a superfamily of enzymes responsible for the metabolism and inactivation of a broad range of toxins [36]. These functions rely on catalysis of the nucleophilic addition of glutathione (GSH) to a wide variety of electrophiles, e.g. toxins, by GSTs. Members of the GST superfamily are extremely diverse in amino acid sequence varying in length from *≈* 210 amino acids in the mammalian organisms up to 300 in plant enzymes. Until 1995 GST was considered to be non-cooperative, but a single residue mutation produced a positively cooperative protein [37]. Later, kinetic studies of *P. falciparum* (protozoan) GST [38] and human GSTs [39, 40] verified allostery be-tween two protein protomers. Negative cooperativity was experimentally found in more recently evolved GSTs, while ancestral GSTs showed a non-cooperative behaviour [41].

For our study we used a crystal structure of GST from *S. japonicum* helminth worm with a non-hydrolyzable GSH analogue S-Hexylglutathione (GTX) [29] (Fig. 1 and Tab.1). This protein-ligand complex simplifies experimental procedures such as isothermal calorimetry, because one does not need to worry about stability of the ligand in a solution. *S. japonicum* GST is a 218 amino acid homodimer with each protomer forming a small N-terminal domain (residues M1–A76) which binds to GSH (LBD), a short linker region (residues D77–G84), and a larger C-terminal domain (residues C85–K218) which mainly composes the substrate binding site (SBD).

There is no published evidence of fluctuation-allostery in GST. Nevertheless, we have chosen to study this protein as a potential candidate for allosteric engineering similarly to the previous mutation study on CAP [12, 42]. Furthermore, holo2 (PDB ID: 1M9A) and apo (PDB ID: 1GTA) X-ray structures produced RMSD 0.52 Å over 216 residue - a small conformational change upon ligand binding (SI Fig. 1) supporting our choice.

### SARS-CoV-2 Main Protease Protein

For the third and final protein used in this study we have chosen a protein related to the SARS-CoV-2 (SCoV2) virus previously, explored for potential drug binding using an ENM approach [14]. The SARS-CoV-2 (SCoV2) main protease (M^pro^, also known as 3C-like protease or 3CL^pro^) (Figure 1). Prior to cleavage, M^pro^ resides on the fifth out of sixteen non-structural proteins of the SCoV2 and is responsible for proteolytic processing of the majority of polyprotein cleavage sites [43]. Thus, it plays a major role in the virus’ life cycle and its inhibition has been a major drug treatment target of the COVID-19 pandemic [44–47]. M^pro^, like CAP and GST, is a homodimeric protein that consists of 306 amino acids. Each protomer can be divided into three domains: domain I (F8–Y101), domain II (K102–P184) and domain III (T201–V303).The substrate binding site is located between domain I and domain II where the Cys-His catalytic dyad performs its function. Domain III is responsible for M^pro^’s dimerization facilitated by a single salt-bridge.

The crystal structure used in this study contains a peptidyl inhibitor N3 (Figure 1 and Tab. 1). This inhibitor binds the substrate binding site and deactivates the catalytic dyad by acting as a Michael acceptor [47]. The preceding coronavirus’ (SCoV) and SCoV2 genetic sequences are approximately 80% identical, reaching 96% amino acid sequence identity between the main proteases [46]. It was experimentally shown that a single N214A mutation dynamically inactivates SCoV M^pro^ [48] while a mutation in the third domain where three consecutive residues were changed to alanie, S284A-T285A-I286A, dynamically enhances the protease’s catalytic activity [49]. The high similarity between the coronoviruses’ proteases and the previously found dynamical control that SCoV M^pro^ exhibits in mutation studies indicates that fluctuation-allostery might be important in the SCoV2 M^pro^, as well.

## Results and Discussion

Both the ENM and all-atom molecular dynamics, including an explicit water model, (aaMD) simulations for the three proteins (see section for details on methods and calculations and SI Fig. 2) generated eigenfrequencies and eigenmode structures for their elastic deformations. From these linear-response measures it is possible to calculate dispersion-relations (frequency of each mode number), the mode-frequency distributions, the B-factor spectra (RMS thermal displacement by residue), and allosteric free-energies of cooperativity.

In the following, these measures are compared between ENMs of different parameter choices against those calculated by molecular dynamics and performed principle component analysis (PCA) on the generated trajectories [50]. Where possible, these parameters were compared to ones measured in experiment. Minimisation of errors gives results, which we report in sequential subsections, for distance cutoff *d*_*c*_, mode cutoff *n*_*c*_ and relative backbone stiffness *ϵ*.

### Distance cutoff

Figure 2 gives the calculated dispersion relation *ω*(*n*) for all three proteins, both in detail for the first 25 modes, then over a much broader base beyond 1000 modes. The information is also given as a distribution function for frequencies (in the third column). In each case, results for ENMs with different choices for distance cutoff *d*_*c*_ (from 7.5 to 15.0 Å, lower cutoffs were discarded due to floppy modes) are compared against the aaMD calculation (in *red*). Immediately apparent is a general pattern by which the ENMs (with generally lower values of *d*_*c*_ a better fit than higher values) capture the fully-atomistic calculations for the first 10 to 15 modes well, then increasingly under-predict the mode frequencies (stiffnesses). A further rapid increase in gradient in the dispersion relation occurs around the 2/3 of all modes in the case of aaMD models only, and in the ENMs beyond the last 100 modes. A consequence of the high-frequency form of the MD dispersion relations is that their frequency distribution is bimodal [26], a feature again not captured by any ENM model (see discussion on the BENM below for both of these points).

**Figure 2:**
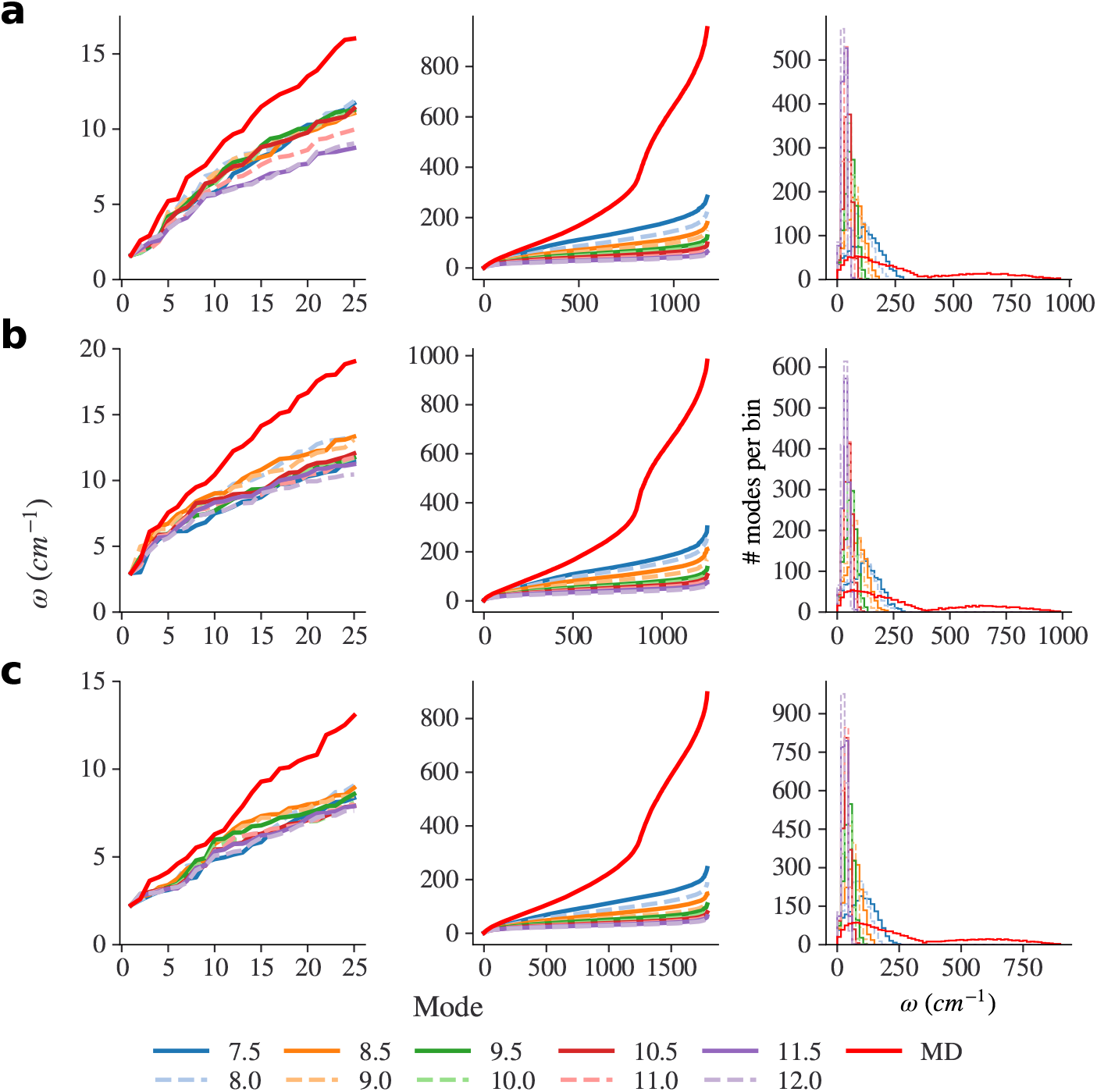
Scanning distance cutoff for the ENM. Each row presents data for a protein: **a** - CAP; **b** - GST; and **c** - M^pro^. First column contains ENM normal mode frequencies for the first 25 non-trivial modes. Second column shown all non-trivial normal modes for a protein. Last, third, column bins all eigenfrequencies in 15 cm^−1^ bins to create density-of-states (DOS) plot.

Figure 3 gives the comparison of B-factors from experiment, aaMD and ENM models of optimal *d*_*c*_ (right column) with the corresponding values of χ^2^ for ENM eigenvalues against aaMD for up to *n*_*c*_=25 normal modes taken in the calculation, and varying *d*_*c*_ (left column). It is striking that the fit for B-factor begins to deteriorate at around the same point in the mode sum that the dispersion relation also deviates (see discussion on mode cutoff below). The same overall pattern against distance cutoff *d*_*c*_ appears as well, with lower (but not the very lowest) values preferred. Table 2 gives B-factor correlation coefficients against X-ray data for the three proteins.

**Figure 3:**
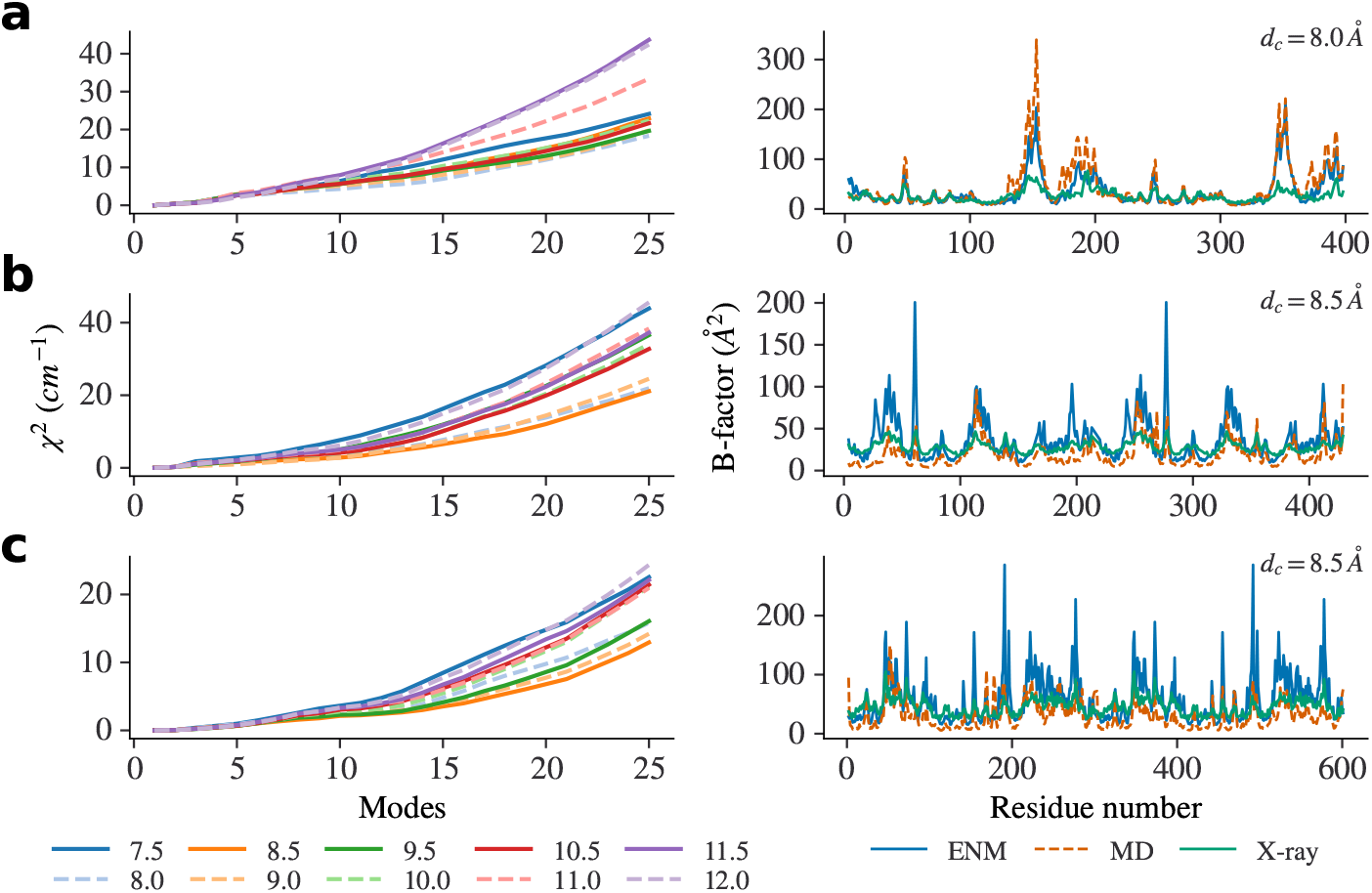
Choosing *d*_*c*_ based on the aaMD simulation results. Each row presents data for a protein: **a** - CAP; **b** - GST; and **c** - M^pro^. (*Left*) χ^2^ for ENM eigenvalues against aaMD for up to *n*_*c*_=25 normal modes. Legend in the bottom labels *d*_*c*_. (*Right*) B-factor plotted against residue number for ENM, molecular dynamics and crystal structure. Values of *d*_*c*_ for each protein are labelled in the same way on the B-factor plot.

**Table 2:** The ENM with varying distance cutoff *d*_*c*_ B-factor correlation with X-ray structural data for apo protein forms. Colouring indicates the correlation magnitude value for each column: small values are coloured in red while large in green. All of the proteins’ non-trivial modes were used for the ENM’s B-factor calculation (See total # modes in table 4).

| $d_c$ (Å) | B-factor correlation | | |
| --- | --- | --- | --- |
|  | CAP | GST | M<sup>pro</sup> |
| 7.5 | 0.748 | 0.781 | 0.615 |
| 8 | 0.736 | 0.792 | 0.716 |
| 8.5 | 0.725 | 0.815 | 0.733 |
| 9 | 0.724 | 0.806 | 0.771 |
| 9.5 | 0.763 | 0.788 | 0.755 |
| 10 | 0.762 | 0.766 | 0.732 |
| 10.5 | 0.781 | 0.750 | 0.731 |
| 11 | 0.761 | 0.738 | 0.754 |
| 11.5 | 0.734 | 0.741 | 0.768 |
| 12 | 0.746 | 0.758 | 0.747 |
| 12.5 | 0.719 | 0.754 | 0.774 |
| 13 | 0.703 | 0.791 | 0.764 |
| 13.5 | 0.684 | 0.783 | 0.775 |
| 14 | 0.686 | 0.781 | 0.787 |
| 14.5 | 0.696 | 0.800 | 0.796 |
| 15 | 0.681 | 0.795 | 0.792 |

The lower end of the distance cutoff range for *d*_*c*_ is set by the requirement that, apart from the first six trivial dynamic modes of translation and rotation, all others are required to be rigid. From the Maxwell-construction, there is a strong constraint from rigidity on connectivity [51]. Calculations of the normal mode frequencies *ω*(*n*) by diagonalisation of the Hessian immediately flag an under-connected ’floppy’ ENM by giving zero-valued eigenvalues beyond the first six.

Focusing first in more detail on the effect of varying *d*_*c*_ on B-factor correlation for non-floppy ENMs, we find that correlation coefficients for all three proteins range between 0.68-0.78, 0.74-0.82 and 0.62-0.8 for CAP, GST and M^pro^, re-spectively (Tab. 2). This range is quite reasonable for a coarse-grained model, and illustrates previous findings that the local differences in B-factors between residues result more from their relative positions in a ’rotational-translational-block’ (RTB) [52] than from spatially highly-localised dynamics. This is because ENM calculations are limited to a low-mode number cutoff so only capture rather non-local dynamics in general (but see discussion of mode cut-off below). Curiously, high correlations can be found both at low and high distance cutoffs for the same protein in all cases, creating a ambiguity of choice for the best *d*_*c*_ value, were the B-factor the only criterion of choice employed.

So motivated, it becomes important to consider the mode structure as well, through the frequency dispersion relation *ω*(*n*). Although this is not directly probed by available experiments, is is possible to make ENM comparisons with aaMD simulations as presented in figure 2. As noted above, for all three proteins shape of the scaled eigenvalues departs faster from aaMD eigenvalues as the *d*_*c*_ values increase. This departure must be a consequence of EN becoming more homogeneous with increasing *d*_*c*_. With a dense network, separate protein domains disappear and protein becomes more comparable to a uniform elastic body, which possesses a dispersion relation of the form *ω*(*n*) *∼ n*^(1/3)^ (in three dimensions) [53]. So, even with the optimal choice of *d*_*c*_ (no zero eigenvalues for non-trivial/floppy modes and EN is not too dense) somewhere around n=10-15, the ENM eigenvalues depart from aaMD results (Fig. 2a) and further separation can be seen in all mode plots (Fig. 2b). Clearly, aaMD high-frequency modes are not represented well by ENMs and are severely underestimated. However, by comparing ENM and aaMD eigenfrequencies using a chi-squared test, it is possible to remove the ambiguity of ENM distance cutoff choice inthe case of each protein.

In figure 3 we present B-factor against residue number calculated by ENMs with appropriate distance cutoff choices based on this double criterion. We have chosen 8 Å cutoff for CAP, as it correlated best with the first 25 modes of CAP’s aaMD simulations as well as being close to the centre of one of the two B-factor optimal ranges. For GST the slightly larger value of 8.50 Å (21.09 cm^−1^) generated a better fit, closely followed by a 8 Å (21.94 cm^−1^) cutoff (with difference smaller than 1 cm^−1^). The best cutoff distance cutoff value for M^pro^, based on our criteria, is also 8.5 Å (12.94 cm^−1^), as for GST, followed by 9 Å (14.21 cm^−1^) and then 8 Å (15.83 cm^−1^).

Using these values, the ENM computed CAP B-factors agree very well both with aaMD and experimental values. B-factor correlation between ENM and experiment is 0.736. In the case of GST and M^pro^, ENMs overestimate some residue fluctuation, more for M^pro^ than GST, but the overall shape is preserved with good ENM to experiment B-factor correlations, 0.815 and 0.733, respectively.

The necessity to choose an optimal value of *d*_*c*_, and the (albeit slight) variation of this value between proteins suggests future comparison against the so-called ’parameter-free ENM’ (pfENM) for computing dynamic allostery [54].

The pfENM does not require *d*_*c*_ parameterization as it employs a continuous decrease of spring constant with distance. It is also prevented from possessing floppy modes due to its fully populated Hessian matrix. However, there is clearly an exchange of parameterisation between the standard ENM and the pfENM as the latter requires a choice of distance-dependent function of spring strength. Nor does pfENM answer the question concerning *n*_*c*_: ”When should we stop summing normal modes?”.

### Mode cutoff

In this section we explore a physical criterion for the appropriate mode cutoff *n*_*c*_, necessary to regularise any sum over modes that is made within the ENM approximation to calculate a physical quantity such as B-factor or allosteric free energies. In the previous section we have already observed empirically how a suitable mode-cutoff might be identified from the dispersion relations. To recap, a universal feature is that ENM spectra *ω*(*n*) follow the MD calculations for the lower (and more spatially distended and correlated) modes, but depart from them at higher mode numbers.

Qualitatively, this is to be expected: the ENM is a coarse-grained model that resolves no spatial structure smaller than the distance between two C-alpha atoms in neighbouring residues. At some point, elastic modes contain structures that possess length-scales comparable to this typical inter-node (inter-residue) distance (SI Fig. 3). For this and higher eigenmodes, the ENM cannot resolve with physical faithfulness the dynamical structure of the real protein, though up to this point the ENM level of coarse-graining ought to be adequate.

The initial surprise, however, from the results presented here systematically, and previously in the case of individual studies, is that the signature of unphysical approximation appears so early in the dispersion relation. In the simple, homogenous elastic case of a spherical solid, the number of modes (three polarisations contained in wavenumber (reciprocal) space within wavenumber *k* grows as *n*(*k*) = (*kR*)^3^/16, where *R* is the radius of the sphere. Since the radii of the proteins in this study are of the order of 10 inter-residue distances, choosing *k* = 10*R* would be a reasonable (homogenous case) estimate for the mode-limit that corresponds to the highest spacial resolution possible, giving a universal estimate for *n*_*c*_ of about 60. Yet in each of the cases examined here, the ENM sum appears to generate unphysical results beyond mode values which (a) vary from protein to protein, and (b) lie between *n*_*c*_ values from 10 to 20 (see figure 2). This is a much harsher criterion than the naive estimate suggested.

The most likely hypothesis for the over-estimation in the naive value is its approximation of uniform elasticity. Much previous work has commented on the universal heterogeneity of the local elastic properties of proteins. Indeed, the viability of the rotation and translation block (RTB) description depends upon precisely that [52], and our previous work, among others, has shown that the mechanism of fluctuation allostery depends on elastic anisotropy, and that the fluctuation-allosteric free energy approaches zero in the case of uniform elastic constant [12]. Material science approaches to the design of allosteric materials have also found that evolutionary algorithms invariably discover solutions of heterogenous elasticity [55].

One consequence of heterogenous elasticity is that the familiar relationship of one wavelength (or equivalently wavenumber) with each frequency of eigenmode, is broken. Each single-frequency elastic mode in a heterogenous medium will exhibit local structure in which the local wavelength varies. Specifically, in regions where the elastic modulus is locally lower than average (corresponding in globular proteins to a low spatial residue density), the local speed of sound is lower than average, and so the local wavelength smaller than average. The criterion for physical resolution of a given elastic mode is therefore much more stringent in a heterogenous material than in a homogenous. For the highest physically-resolved mode is now the one whose smallest local wave-length (or inter-node distance) interferes with the spatial resolution of the coarse-grained model, not the average wavelength.

We tested this hypothesis by a semi-quantitative approach. We visually represent the normal mode structures for each protein around the values of *n*_*c*_ suggested by the dispersion relation modelling. Visual inspection of the eigenvector structure identify regions where closely located EN eigenmode nodes have large negative eigenvector product. In the case of each protein, the locally ’soft’ region which is responsible for the failure of spatial resolution is identified. Figures 4, 5 and 6 present for the cases of CAP, GST and M^pro^ respectively, the structure of three eigenmodes, as wall as identifications, and close-up visualisations, of the regions of first-failure. As a quantitative metric, we use a simple estimate of two vectors’ velocity cross-correlation factoring in proximity of two nodes:

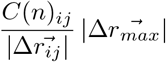

where *C*_*ij*_ is cross-correlation of dynamic for a given mode(See section) and 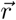 is a distance vector between two nodes. The separation distance between two nodes is normalised in respect to the largest inter-node distance in the ENM. More negative values troubleshoot regions where nodal surfaces become closely located.

**Figure 4:**
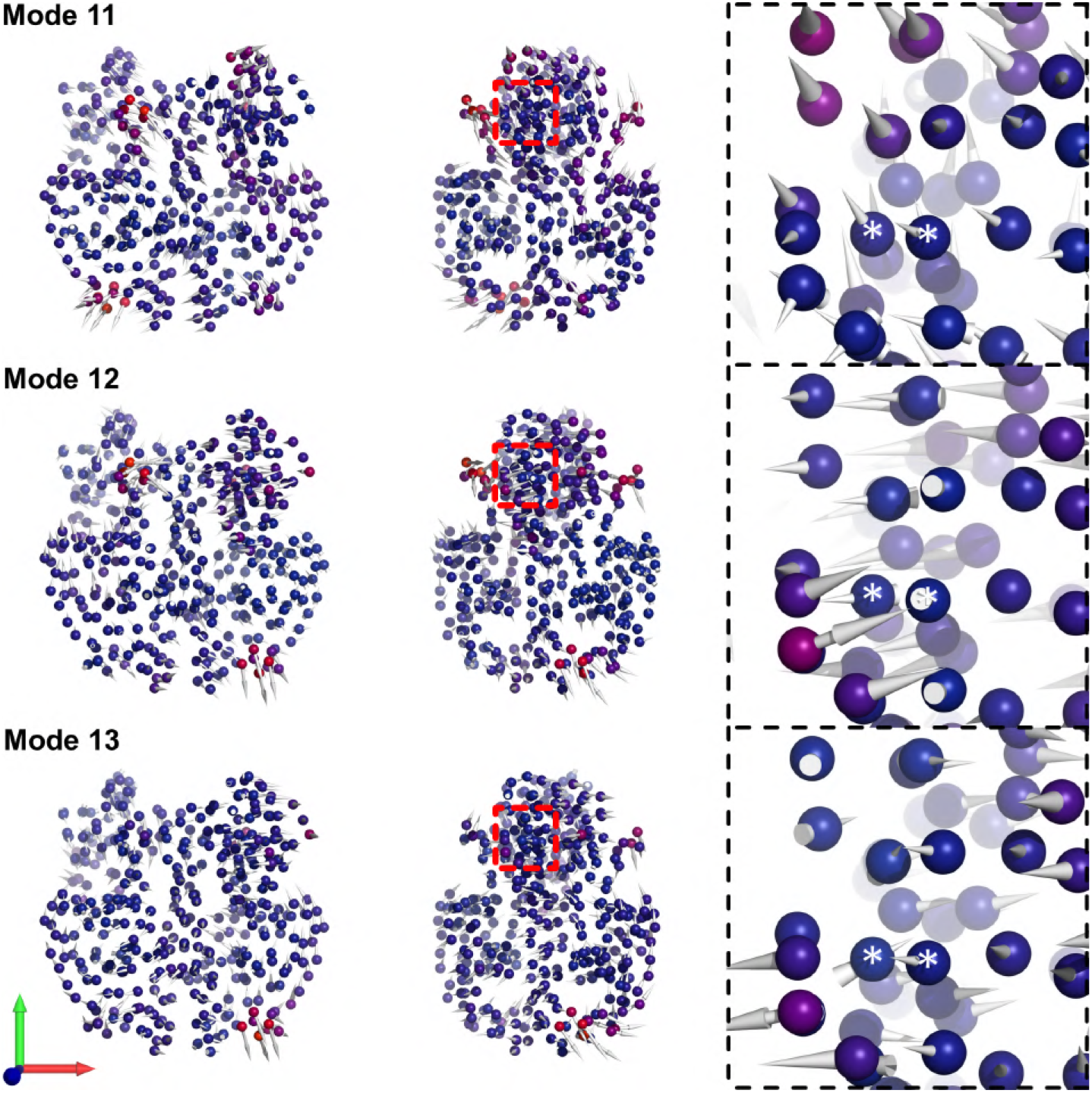
CAP’s ENM (*d*_*c*_ = 8 Å) normal modes 11, 12 and 13. (*Left*) Front view of a mode. Spheres represent EN nodes and arrows mode’s eigenvectors. Nodes are coloured by normalised eigenvector’s RMS magnitude: blue nodes have small values near 0, while red ones towards 1. (*Middle*) Side view (− 45° rotation around y-axis). The red squares show a problematic region for the ENM with a rapid spatial gradient of eigenvector. (*Right*) The problematic region (CAP’s C terminal region). Nodes that are close to nodal surface are marked with an ”*”. Our eigenvector change metric between the two neighbouring nodes produced 16.22, −10.32, −6.33 for modes 11, 12 and 13, respectively. Left-handed axis in the bottom left corner x, y and z axis coloured in *red, green*, and *blue*, respectively.

**Figure 5:**
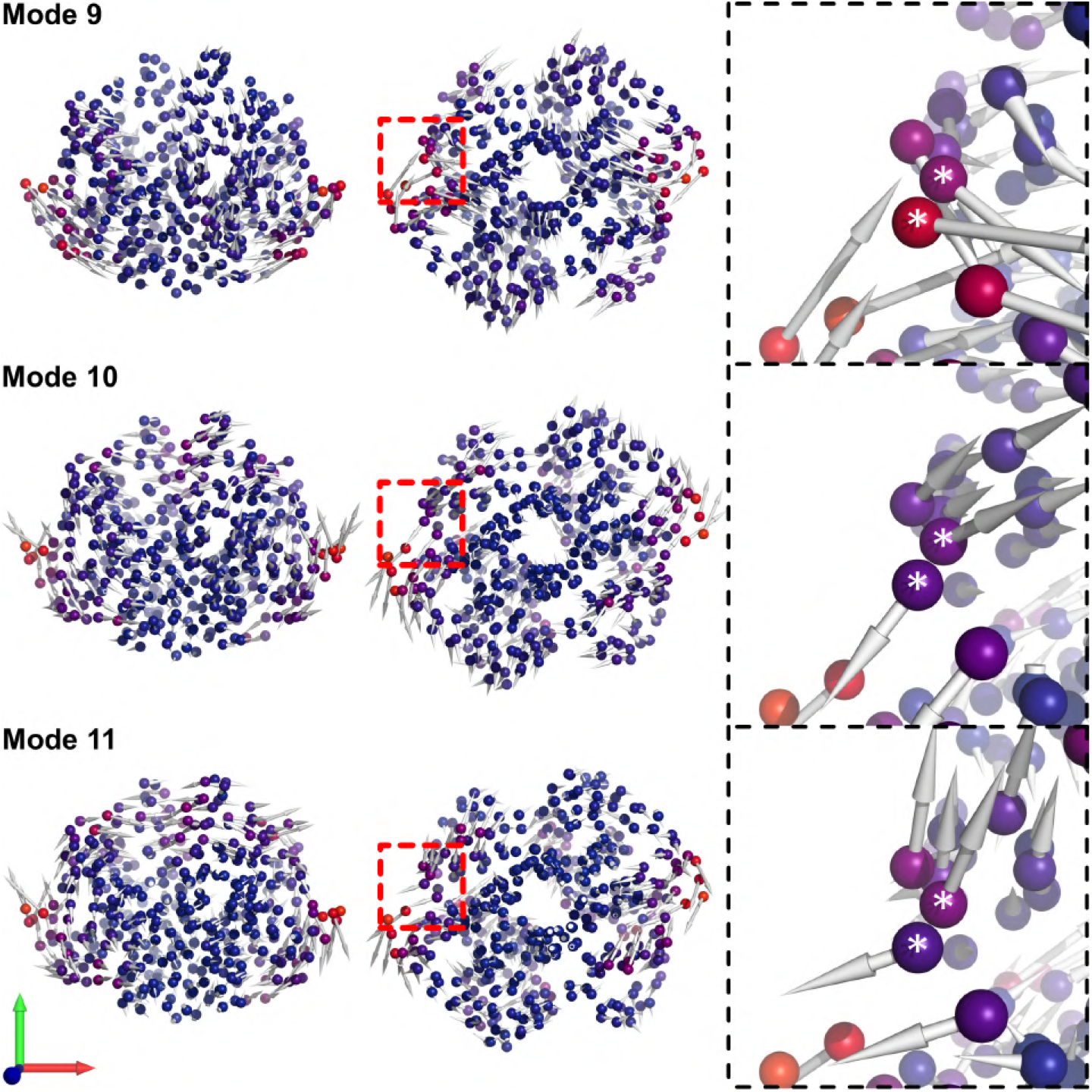
GST’s ENM (*d*_*c*_ = 8.5 Å) normal modes 9, 10 and 11. (*Left*) Front view of a mode. (*Middle*) Side view (−90° rotation around x-axis). (*Right*) The problematic region (GST’s side loop region). Our eigenvector change metric between the two neighbouring nodes produced 14.11, −8.99, −9.87 for modes 9, 10 and 11, respectively.

**Figure 6:**
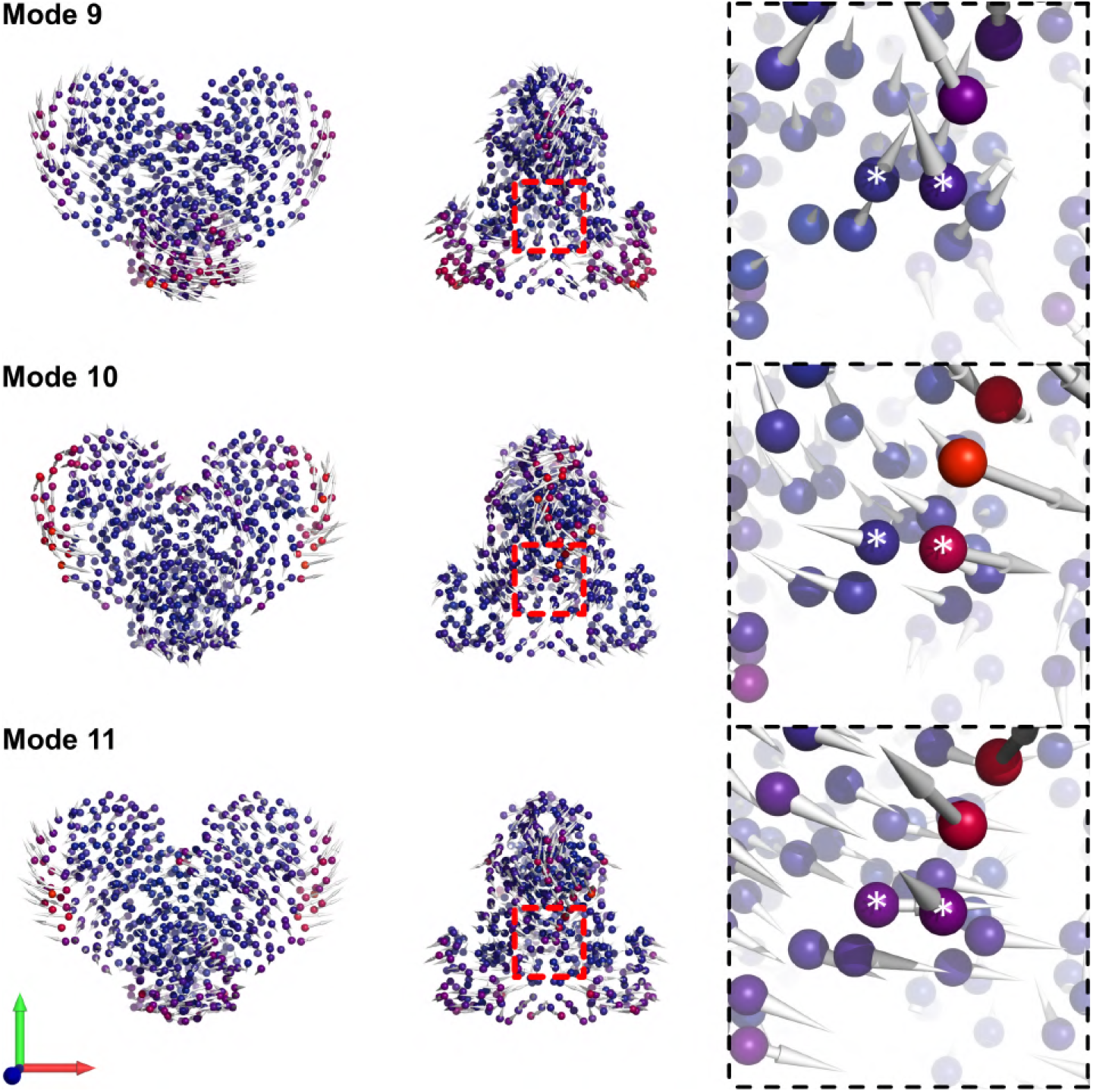
SCoV2 M^pro^’s ENM (*d*_*c*_ = 8.5 Å) normal modes 9, 10 and 11. (*Left*) Front view of a mode. (*Middle*) Side view (90° rotation around y-axis). (*Right*) The problematic region (M^pro^’s side loop and a small helix regions). Our eigenvector change metric between the two neighbouring nodes produced 15.44, −19.31, −6.97 for modes 9, 10 and 11, respectively.

Investigation of CAP’s normal modes did not show obvious neighbouring eigenvector change for the first 25 modes, compared to GST and M^pro^ (See Fig. 5 and 6). However, modes 11, 12 and 13 indicate a locally soft region near the homodimer’s DBD where the local direction of displacement between neighbouring residues becomes orthogonal and reverses (See Fig. 4 highlighted in *green*). The region undergoes a rotational motion, with axis of rotation passing through the DBD. Comparing this with the dispersion relation indicates that this is the first unphysical structure counted in the mode sum. Of course at this point the failure is physically constrained to one small region of the protein, but beyond this point the model sum will incorporate an accumulation of such local failures.

In the case of GST, the departure of the ENM calculation from that of aaMD occurs slightly earlier in the mode sum. So figure 5 presents eigenmode structures in the same form as above, for CAP, but in the mode range 9, 10 and 11. The first problematic region in the case is identified in a side-loop, which is another likely secondary structure of locally low residue-density.

A similar mode-region displays the characteristic divergence in the case of M^pro^. Figure 4 shows the problematic region for modes 9, 10 and 11. The rapid and poorly-resolved mode structure appears in a region containing a side-loop and small helix, first identified in mode 10.

To summarise, a departure of the ENM eigenvalues from aaMD PCA eigenvalues occurs at a different mode number for each of the proteins: CAP 11^th^-13^th^, GST 10^th^-12^th^ and SCoV2 M^pro^ 10^th^-12^th^ modes, as well - but no later than 25^th^ mode, which we chose for chi-square evaluation. In each case, clues in eigenvector structure appeared around those modes where deviation from the aaMD results set in. This confirms the hypothesis that a mode becomes problematic for a given EN resolution when a rapid change of velocity happens between neighbouring nodes.

All highlighted regions that pose problems for the model are located on the proteins’ surface, where EN nodes are less connected. This sparse connection means higher flexibility, especially for loops or terminal regions, like in CAP’s N-terminal helix case. We should recall here, as a methodological caveat, that in extreme cases, such local under-connectivity of nodes can cause floppy (zero eigenvalue) modes. This is avoided by trimming protein termini before ENM simulations. The work here indicates that even when floppy modes are removed in this way, the problem of locally poorly-connected regions can still become problematic in an early failure of physicality of the mode sum.

### BENM

The final question of our list to explore is that of the consequences for modelling protein dynamics and allostery of endowing special (stiffer) values of inter-C-alpha bonds to those residues adjacent within the protein backbone, recognising the greater bond-strength expected from their covalent connectivity when compared with off-backbone bonds within the globular structure.

To explore the effect of backbone enhancement, all calculations were performed on ENMs of the three protein structures generated with *d*_*c*_ = 8.0 Å or 8.5 Å and a range of values for *ϵ* from 1 (standard ENM) to 200. Figure 7 illustrates the form of the resultant ENMs for each of the three subject proteins, using thicker lines to represent the embedded backbone path of stiffer bonds, and to visually encode their relative stiffness.

**Figure 7:**
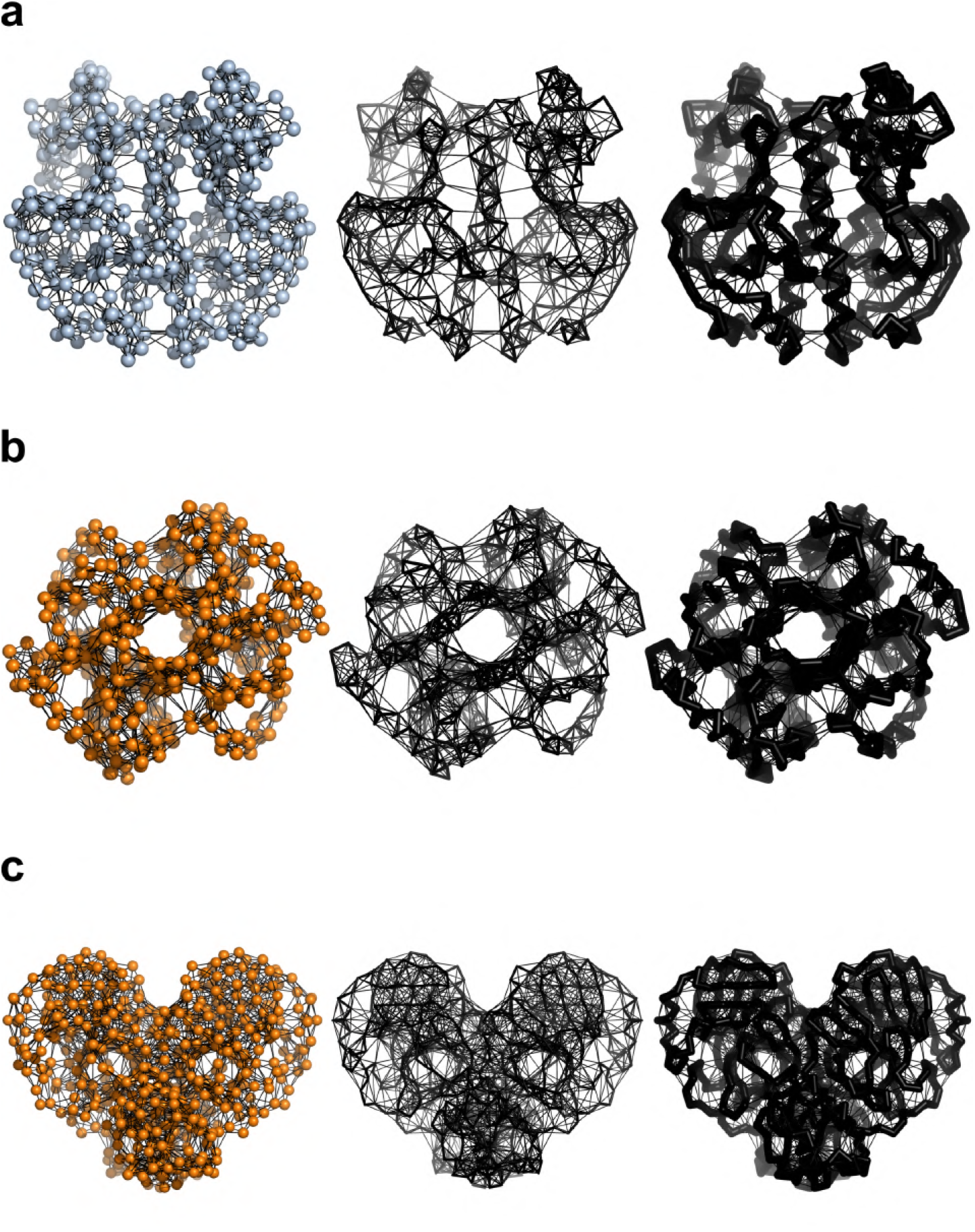
Apo protein forms of the BENM for *ϵ* = [1, 10, 100]. Each row presents data for a protein: **a** - CAP; **b** - GST; and **c** - M^pro^. Elastic network nodes are shown only for BENM with *ϵ* = 1 for clarity. Colour of EN nodes corresponds to the colour form the distance cutoff scan. Backbone springs’ radius is calculated as 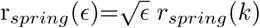 equals EN spring radius for uniform spring constant.

Leaving aside the question of allosteric cooperativity for the following section, here we focus on the effect of backbone stiffening on the frequency dispersion and distribution. The calculations are presented in Figure 8, with the main qualitative result that BE results in a new, high frequency, peak to the mode distribution (as previously found by [26]) in all cases, and ostensibly agreeing qualitatively with aaMD results. Table 3 captures the renormalisation that the BENM models require on the baseline spring constant (tuned to match the lowest-frequency aaMD mode frequency), and table 4 summarises the structure of the mode distributions in terms of the ratio of the two peak frequencies.

**Figure 8:**
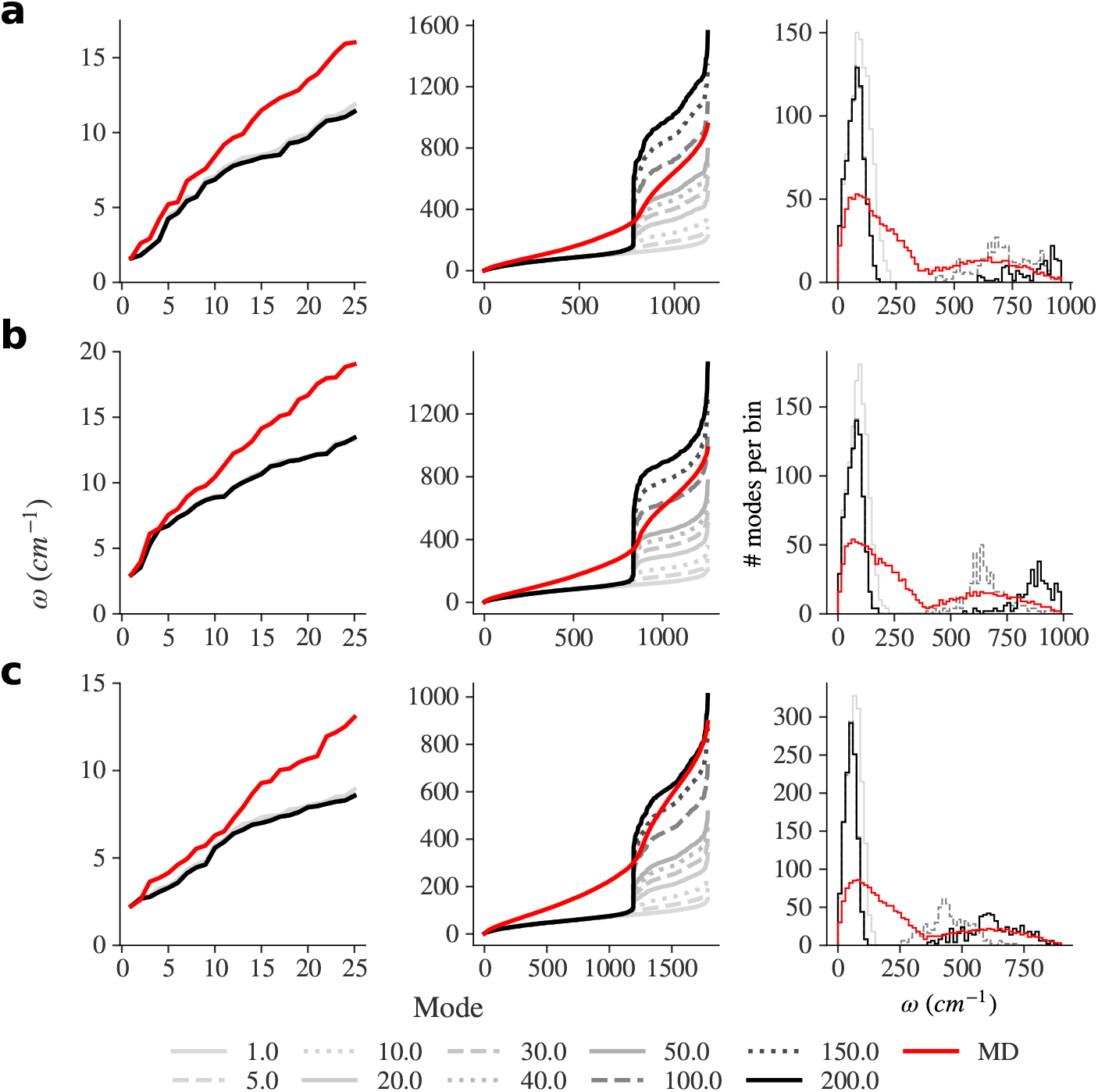
BENM scan results for the mode frequency dispersion relation (left column, low modes only, centre column all normal modes for each protein modes), and frequency distributions (right column). Molecular dynamics results in red, and the range of values for *ϵ* given by grey-scale with alternating solid, dash-dash and dotted lines. For each DOS plot *ϵ* = 1, 100 and 200 are shown, and the bin width equals 15 cm^−1^.

**Table 3:** Tuning BENM. ENM uniform spring constant *k* scaling for proteins’ BENMs with increasing backbone enhancement coefficient *ϵ* and χ^2^ test results for DOS of the BENMs and aaMD results. BENM *ϵ* = 1 corresponds to the classical ENM without backbone coefficient enhancement. *k*_*scaled*_ is tuned based on the aaMD first mode frequency. DOS bin width equals 15 cm^−1^. χ^2^ in bold are the point at the optimal *ϵ* for the given BENM. Distance cutoffs are 8, 8.5 and 8.5 Å for CAP, GST and M^pro^, respectively.

| | Protein | $\epsilon$ | | | | | | | | | |
| --- | --- | --- | --- | --- | --- | --- | --- | --- | --- | --- | --- |
|  |  | 1 | 5 | 10 | 20 | 30 | 40 | 50 | 100 | 150 | 200 |
| $k_{scaled}$<br>(kcal mol<sup>-1</sup> Å<sup>2</sup>) | CAP | 31.10 | 25.61 | 24.60 | 24.04 | 23.84 | 23.74 | 23.68 | 23.56 | 23.52 | 23.50 |
|  | GST | 25.71 | 22.09 | 21.35 | 20.92 | 20.77 | 20.70 | 20.65 | 20.55 | 20.52 | 20.50 |
|  | M<sup>pro</sup> | 12.28 | 9.99 | 9.51 | 9.24 | 9.15 | 9.10 | 9.07 | 9.01 | 8.99 | 8.97 |
| $\chi^2$<br>(cm<sup>-1</sup>) | CAP | 1406 | 961 | 887 | 1407 | 1668 | 1340 | 1166 | **893** | 1167 | 1180 |
|  | GST | 1932 | 1370 | 1254 | 1486 | 2527 | 3256 | 2209 | **1171** | 1314 | 1789 |
|  | M<sup>pro</sup> | 3723 | 2631 | 2338 | 2322 | 2417 | 2729 | 2976 | 2893 | 2435 | **2348** |

**Table 4:** The BENM mode frequency comparison for apo protein forms. Number of the total non-trivial modes (3*N* − 6), number of low-frequency and high-frequency normal modes in the BENM’s bimodal distribution, and ratio between the two distribution’s modes.

| Protein | Modes (#) | $\omega_{low}$ (#) | $\omega_{high}$ (#) | $\omega_{low} : \omega_{high}$ |
| --- | --- | --- | --- | --- |
| CAP | 1176 | 784 | 392 | 2:1 |
| GST | 1254 | 837 | 417 | 2.01:1 |
| M<sup>pro</sup> | 1788 | 1193 | 595 | 2.01:1 |

The first new detailed result emerges, however, not from the distribution of mode frequencies, but from the dispersion relation. Although backbone enhancement does indeed reshape these curves which now match aaMD results better, the improvement is only seen in the higher-frequency mode region. Specifically, the BENM results in the emergence, in each case, of a critical mode number, below which there is no modification to the dispersion curve (and so mode frequencies) at all, and at which the introduction of BE produces an abrupt step to a higher-frequency branch of the curve at around the mode ranked 2/3 of all normal modes in increasing frequency, for each protein. So, well-tuned BENMs manage to capture the eigenvalues right after this transition. Higher choices of *ϵ* are able to optimise fit at the highest frequencies, but this is not likely to be a physical choice, and it is not possible to match the regions of the dispersion curve at the step, and at higher frequencies as well, with a single value of *ϵ*. The previous BENM study by Ming and Wall [26] present much better agreement between ENM, BENM and aaMD density-of-states (DOS) (albeit for a different protein) than the model presented here. However, we note that they used normal mode analysis (NMA) instead of PCA, as in this work, as a comparison for aaMD data. The ENM can be viewed as a coarse-grained version of NMA since both explore one local energy minimum set by the initial protein structure, while PCA uses data from aaMD simulation that has an ability to access different energy minima. This inability to capture behaviour of the high frequency region except at the discontinuity suggests that at least another parameter must be included into the ENM to capture broad regions of the high-frequency modes. Comparison of the BENM dispersion relations with those from aaMD strongly suggests that the principle difference in the physics of the two is the presence of a continuous range of effective spring constants in the atomistic model. It is clearly the discrete bimodality of spring constants in the BENM models that gives the two discrete regions in their dispersion curves, and the discontinuous step in frequency between them. Recent work that optimises ENM models to aaMD by permitting just such a continuous parameterisation of bond strengths supports such a conclusion [21]. We may also deduce the interesting result that all dynamic modes up to the critical mode number of the BENM discontinuity operate without significant stretching of the protein backbone.

Varying the backbone enhancement shows that greater *ϵ* values always affect the newly-generated high-frequency region by increasing its values. There is a larger backbone enhancement to frequencies in the region of the eigenvalue discontinuity. To add some quantitative detail to this effect, we observe, at the discontinuity, a low to high frequency ratio of 2:1 for all three proteins (Tab. 4), when the ENMs are best-matched to aaMD simulations. The same ratio was reported in the BENM work on bovine trypsinogen [26]. Our findings therefore support work by authors of the latter study that this ratio might be a universal feature of globular proteins’ BENM. However, we now see more clearly how the apparently much-improved distributions are achieved: this is through a rather crude approximation to a high-frequency branch of the dispersion curve. However, the result of this work on the optimum value of *ϵ* is more delicate: table 3 suggests a value of 100 for both CAP and GST, but the dispersion relation for M^pro^ is optimised at the transition region better using a value of 200. A wider study of other proteins would be helpful in resolving how universal the BENM is able to be.The same non-universality emerges, however, if the BENM is matched only at the discontinuity, rather than over the entire higher-frequency branch. Values for *ϵ* = 50 match aaMD values well for about 50 modes after the critical mode for CAP and GST, but about 100 in this case for M^pro^. As we will find, BENM has no effect on proteins’ cooperativity.

### Cooperativity

Finally, we assess the effect of distance- and mode-cutoffs, and of backbone stiffening, on the allosteric co-operativity of effector binding. The fundamental effect here is a difference in free energy difference between the holo2/holo1 binding event, and the holo1/apo binding event. Figure 10 displays the effect of the joint cut-offs (left column) and on backbone stiffening (right column) for all three test proteins.

We note that, for allosteric response, there is a range of choices by which the effector binding structure is modelled (Fig. 9). The simplest case is to represent ligand as a single node with the corresponding ligand atomic mass, as shown in the figure 9 middle panels. A more detailed ligand model accounts for each heavy-atom (non-hydrogen atoms) as a EN bead assigned its own atomic mass, as seen in the same figure left panels. If ligand EN beads are treated in the same way as protein EN beads then the number of protein-ligand bonds will grow as the distance cutoff is increased. However, the importance of ligand connectivity to protein chain in ENM has been previously found paramount for estimating cooperativity in CAP [42]. For example, in an ENM of the CAP where cAMP is represented as a single node and treated in the same way as the protomer nodes, the number of springs attached to the ligand grows the larger the distance cut-off. The 8.0 Å ENM receives three additional springs between the cAMP node and neighbouring residues, compared to the ENM using a value of 7.5 Å, two of which connect oppositely to the first, on the other protomer (SI Fig. 4). In order to respect the crucial geometry of biochemical binding of the ligands, we use a custom protein-ligand bonding for each protein based on biochemical information obtained from the PoseView software tool [56] and PDB database [57] (SI Fig. 5). We have chosen the single-bead ligand model to present cooperativity results because it is the simplest biophysical model that takes into account the biochemistry of active site and, as we will see, fully-atomistic treatment of ligands will not change the results qualitatively. Moreover, our results show that if ligand beads are not treated specially in this way, but simply as the rest of the protein beads, i.e. ligand beads have an isotropic distance cutoff, *K*_2_/*K*_1_ is non-cooperative for larger distance cutoffs (SI Fig. 6).

**Figure 9:**
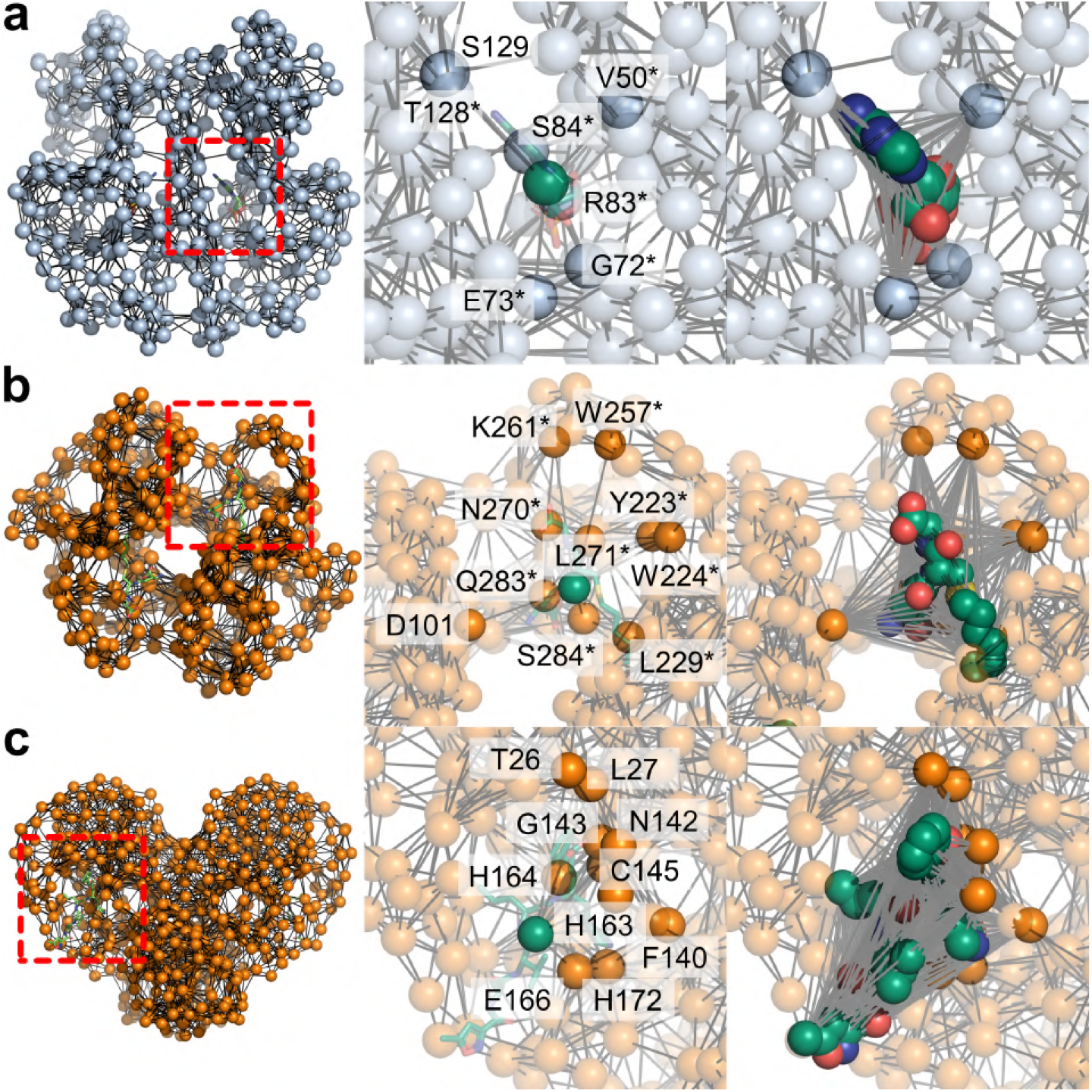
Cooperativity for ENM distance cutoff and BENM scan. The dashed rectangles on the plots represent non-cooperative region. Each row presents data for a protein: **a** - CAP (*d*_*c*_ = 8 Å); **b** - GST (*d*_*c*_ = 8.5 Å); and **c** - M^pro^ (*d*_*c*_ = 8.5 Å). *Left* shows an ENM of the protein with stick representation of a ligand, *middle* presents a ligand modelled as a single mass-weighted with biochemically relevant bonds (EN springs, see labelled residues) and *right* shows all heavy-atom EN ligand model with the same protein-ligand bonding rule as for the protein itself. In the single bead ligand model, the ligand node is assigned the full ligand mass (see Tab. 1), while the in the heavy-atom ligand model each node has corresponding atomic mass. In the latter model, each ligand’s EN node is connected to the specified protein node and all ligand nodes are interconnected with each other. In *left* subfigures red squares show enlarged active site regions. Asterisks mark residues from protomer B.

**Figure 10:**
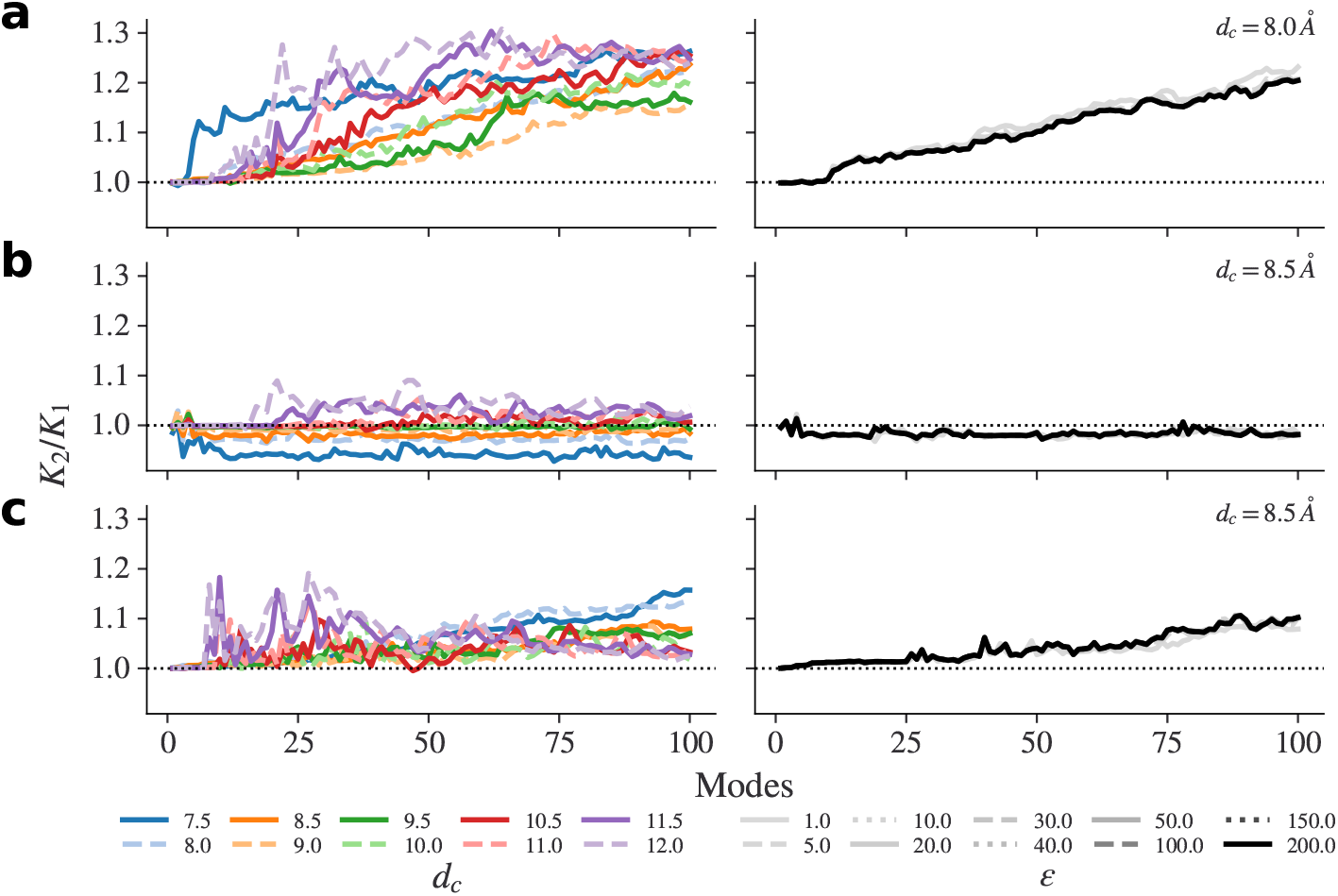
Cooperativity for ENM distance cutoff and BENM scan. Dotted line on the plots represent non-cooperative region. Each row presents data for a protein: **a** - CAP; **b** - GST; and **c** - M^pro^. *Left* shows results for distance cutoff scan while *right* presents backbone-enhancement scan results. Each sub-plot represents the calculated cooperativity values against the total number of summed modes. Only the first 100 non-trivial modes are shown. In each case, the ligand was modelled as a single mass-weighted elastic network bead with a total mass of the ligand and the protein-ligand interaction set to one 9.

All three chosen ENMs show behaviour expected from previous studies: CAP and M^pro^ are negatively cooperatively, while GST is non-cooperative. From previous studies [42], this is likely to be because a single-node approach ENM does not take into account the structural biochemistry of the ligands (treating the ligand node as equivalent to a alpha-carbon atom, although with the right total ligand atomic mass). In the models studies here, ENM’s protein-ligand connectivity is determined by initial ligand heavy atom placement and chosen distance cutoff only.

For all values of distance cutoff apart from the smallest investigated, the ENM predictions for CAP stay on the non-cooperative line until mode 10-15, after which the negative cooperativity starts to grow. For CAP ENMs with *d*_*c*_ =7.5 Å cooperativity suddenly becomes negative in course of three modes, from 4^th^ to 6^th^ mode and keeps gradually increasing afterwards. All ENMs based on GST’s crystal structure produced nearly non-cooperative binding predictions over the presented range of distance cutoffs. Again, as in CAP’s case, only the lowest *d*_*c*_ makes a slightly positively cooperative ENM. For M^pro^ all ENMs stay non-cooperative for the first 7 non-trivial modes, afterwards the cooperativity fluctuates with mode cut-off, but remaining consistently in the negatively cooperative region. In all three cases, the numerical results for cooperativity fluctuates in value as a function of mode cutoff below 50^th^. However, beyond this point, the fluctuations reduce in amplitude and converge towards physically acceptable value of cooperativity. This suggests that even beyond the mode number where the strict uniform criterion for physical resolution is first broken, yet higher modes still carry information on the cooperativity.

We observed another delicate balancing feature of mode and distance cutoffs in regard to cooperativity: ENMs with lower distance cutoffs approached their expected cooperativity values earlier in the mode summation, but overestimate the value as we keep adding more modes. On contrary, higher cutoffs converge to the expected value very well, but require more normal modes than presented here. When all heavy-atoms were included in the ligand modelling, the calculated cooperativity is of the same magnitude and sign as for the single-bead ligand model (SI Fig. 7). For the higher *d*_*c*_ we see less fluctuation for the first 100 non-trivial modes than in a single-bead ligand model case.

Significantly, little or no change in cooperativity is seen for low-frequency modes when BENM is applied to the proteins. This small effect of BENM on the numerical cooperativity is in accord with the findings of the previous section: only higher frequency modes are noticeably affected by the backbone enhancement. Once this is combined with the complementary finding in the literature that allostery is principally communicated by global, and hence low-frequency, modes, the result is that allostery is expected to be unaffected by backbone enhancement.

## Conclusions

The exhaustive study of three homodimeric proteins, each offering the in-principle possibility of fluctuation-allosteric signalling between the effector binding pockets, has provided an extensive test of the ENM model in regard to the dynamics, and the thermodynamics, of binding.

The first finding, in accord with previous studies, is that although the ENM is coarse-grained at the level of entire residues, it is able to give good accounts of the B-factor, and low-frequency mode-dispersion measures found both experimentally and in fully-atomistic simulations. This confirms the insight that even local differences in B-factor arise principally from the rotations and translations of large units within the globular protein.

Second, the choice of distance cut-off between bound and unbound residues within the ENM is optimised within rather tight limits. If too low a value is chosen, the ENM becomes under-constrained and under-rigid in the Maxwell sense. Once a sufficient number of local bonds is present, the eigenmode dispersion relation (of mode frequency with mode number) becomes a sensitive quality-measure of the model, when compared to fully-atomistic simulations. Too great a cutoff distance creates, in every case, too sharp a ’roll-off’ in the dispersion relation when compared with the aaMD results. When the minimum error criteria for dispersion curve and B-factors are combined, all three structures investigated suggested optimal models with cutoff distances at or just above a choice of 8.0 Å. The reducing effect of a longer cutoff distance on the co-operativity is consistent with the notion that adding more distant bonds artificially dilutes the effect of a few that are key for allostery. The denser structures are also, in consequence, more elastically homogenous, which is known to decrease the effect of fluctuation allostery [42]. However, the principle conclusion in regard to the allostric cooperativity of ENM models is that, as previously found in one case [42], to capture the correct physics it is necessary to resolve the ligand structure more carefully than that of the surrounding protein, especially in regard to those residues of the host protein to which it makes significant bonds.

Third, the way that bound ligands are modelled in conjunction with ENMs is important in capturing numerical predictions for cooperativity, especially for higher distance cutoffs. In particular, there is no requirement that the ligand, which plays a special, local, role in allostery, be modelled in the same way as the rest of the protein. In particular, we found that a single-bead ligand representation model does not capture allostery adequately, including poor convergence, if the bead is treated as protein EN beads. This is due to the generation of unrealistic ligand-protein bonds. However, when the ligand is connected only to the biochemically relevant residues around the active site, calculated cooperativity is consistently converging over the range of distance cutoffs used for the protein EN beads. No qualitative difference has been seen between single-bead and many bead ligand representations if the ligand is connected to only biochemically relevant residues around the active site for the chosen proteins. However, The single-bead ligand model does not, of course, capture the correct aspect ratio of the ligand molecule when it is of an extended form, and might not necessarily have correct bond angles with the connected residues in comparison to the all heavy-atom model, but this model is less computationally demanding and easier to construct. These findings will motivate further research in ligand modelling beyond the simple Hookean potential for protein-ligand elastic network springs and coarse graining of its structure to capture correct dimensional ratio of the ligand molecule.

Fourth, the enhancement of the protein backbone stiffness, which is possible within ENM model (to give the BENM family of models) has the most striking effect on the distribution function for eigenmode frequencies. As first found by Ming and Wall [26], the BENM typically possesses a bimodal distribution. However, analyses of the full dispersion curve show that backbone enhancement has a much stronger effect on modes typically higher than the 800th normal mode (or generally, at 2/3 of the ordered modes), than on lower modes, whose effective stiffnesses are only mildly perturbed. It should be noted that at this point, the corresponding eigenfunctions are not generally of physical significance at the level of coarse-graining associated with the ENM. Furthermore, the resulting discontinuity in the frequency dispersion, not present in aaMD calculations, indicates that more quantitatively faithful ENM models need to respect a continuous range of inter-residue bond strengths, as found in [21].In no case, however, was the allosteric free energy and co-operativley found to be affected by the backbone stiffening. Notwithstanding, a backbone stiffness ratio of 100 does give a better match of the dispersion spectrum in the narrow range of modes clearly dependent in a dominant way on that property, close to the discontinuity in the frequency dispersion.

Fifth, the heterogenity of effective elasticity in all proteins results in a much sharper criterion for the ability of coarse-grained models to resolve the finest structure in even the lowest, and softest (and so often most global) of dynamic modes. This is because softer regions contain much shorter local effective wavelengths for dynamic modes than their structure in stiffer regions. Therefore, the mode number at which unphysical effects, due to poor local spatial resolution, begin to appear, can be much lower than that expected for homogenous elastic solids similarly coarse-grained. It also means that there is no universally-appropriate mode-cutoff for ENMs across all proteins. In the case of the three proteins studied here, unphysically small-scale local structures appeared between the 10^th^ and 15^th^ mode. This is at least a factor of four lower in the mode-ordering than would be the case for a similar body of homogenous elasticity, yet could be identified and correlated with the point at which the ENM dispersion curve departed from that calculated from molecular dynamics, confirming the hypothesis that the region of first failure of coarse-graining in regard to the fine-detail of eigenmode structure determines the range-limit of physical ENM modes.

Elastic Network Models have proved themselves to be immensely powerful tools for exploring protein dynamics efficiently at the larger lengthscales of protein physics. However, they turn out to be more subtle that a first glance would suggest, and to demand a nuanced and careful treatment, including the parameterisation and choice of their necessary cutoffs, if they are to speak to experimental data adequately.

## Materials and Methods

### Molecular Dynamics simulations

The molecular dynamics (MD) simulations employed the ff19SB force field within the AMBER 17 simulation program for the proteins [58]. ff19SB force field is used as the energetic interactions of side chains, which outperforms the ff99SB and ff14SB force fields [59]. MD calculations used a short-range cut-off of 10 Å, with the long-range portion of the Coulomb potential represented by an Ewald summation, and employed a time step of 2 fs. The bond lengths were constrained by the SHAKE algorithm. The starting structures were obtained directly from Protein Databank (PDB) X-ray diffraction structure [57]. `pdb4amber` program performed default protonation for the side chains while the force field, water molecules and ions were added to each system using `tleap` program, both programs from AmberTools21 package [60]. The water molecules were represented by the “optimal” 3-charge, 4-point rigid water model (OPC) force field [61], which is a recommended water model for ff19SB force field [59]. Periodic boundary conditions were employed using a periodic truncated octahedron. The net charge of the proteins was not neutral, so either sodium or chlorine ions were added to neutralise the charge. The systems’ energy was minimised in two stages; firstly protein was held with restraints while the water was energy minimised, then the restraints were removed and the entire system energy minimised. The temperature of each system was then raised to 300 K over a 20 ps period and density equilibration was performed over the next 20 ps by switching on pressure coupling using the thermostat and barostat. The systems were then simulated for up to 500 ns, with up to 200 ns of the simulation being kept as the equilibration time. Principal component analysis (PCA) was performed by diagonalising the mass weighted covariance matrix of the atomistic simulations using pytraj program from AmberTools21 [60]. The eigenvectors represent the shape of the atomistic motion and the corresponding eigenvalues the extent of the motion. As the each system differs, the simulations settings are included in table 5.

**Table 5:**
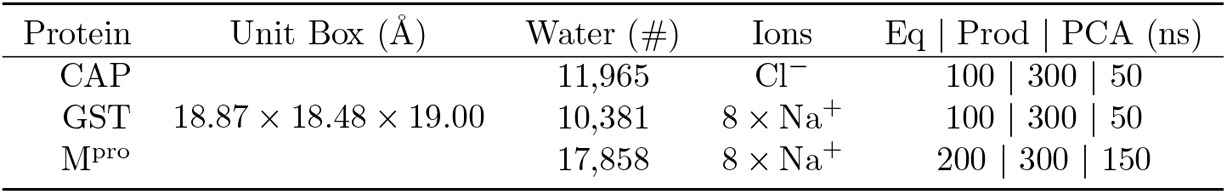
MD simulation parameters. Truncated octahedron PBC unit box size; number of water molecules: ions to neutralize protein’s charge (water molecules were replaced by added ions); and system’s equilibration, production times with the time slice (form the end of production) used for PCA.

| Protein | Unit Box (Å) | Water (#) | Ions | Eq \| Prod \| PCA (ns) |
| --- | --- | --- | --- | --- |
| CAP |  | 11,965 | Cl<sup>-</sup> | 100 \| 300 \| 50 |
| GST | 18.87 × 18.48 × 19.00 | 10,381 | 8 × Na<sup>+</sup> | 100 \| 300 \| 50 |
| M<sup>Pro</sup> |  | 17,858 | 8 × Na<sup>+</sup> | 200 \| 300 \| 150 |

### Coarse-grained simulations

Normal Mode Analysis (NMA) of ENM describes protein motions around equilibrium and can be used to calculate the partition function for large scale harmonic thermal fluctuations in protein structure, including those responsible for allostery [62]. Two main approximations of NMA are:

- The structure fluctuates about a local energy minimum. Consequently no other structures beyond the given equilibrium can be explored.
- The force field everywhere arises from sums over ENM harmonic (spring) potentials.

The whole NMA method can be reduced to three steps:

1. Construct mass-weighted Hessian for a system. For a protein ENM the system consists of the co-ordinates of the alpha-carbon atoms (*N*) for each residue from the corresponding PDB structure.
2. Diagonalise the mass-weighted Hessian to find eigenvectors and eigenvalues of the normal modes.
3. Calculate the partition function (and so free energy) from the product over the normal mode harmonic oscillations.

The diagonalisation of the 3*N* × 3*N* mass-weighted Hessian matrix is written as

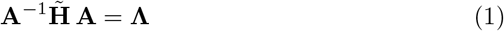

where 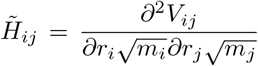 the potential energy function *V*; distance be-tween nodes *r*; node masses *m*. The eigenvectors of the mass-weighted Hessian matrix, columns of **A**, are the normal mode eigenvectors **a**.

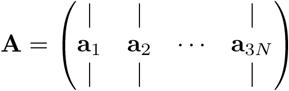

**Λ** is a 3*N*×3*N* diagonal matrix with diagonal values equal to the associated normal modes’ (3*N* − 6 non-trivial) squared angular frequencies *ω*^2^. The potential function used in this study is:

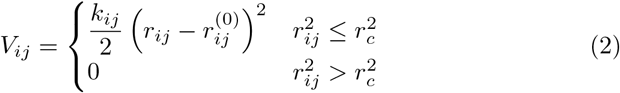

where *r*_*c*_ is a cut-off radius; while *r*^(0)^ is the equilibrium distance between nodes derived form PDB crystallographic structure. For the wild-type protein, all spring constants (also sometimes denoted as *γ*) are equal *k*_*ij*_ = *k* = 1 kcal Å^−2^ mol^−1^.

For the given interaction network, the eigenvectors **a** are independent of spring constant k. Thus, *ω*^2^ ∝ *k* which allows us to scale eigenfrequencies without reevaluation of the ENM. To match the first mode eigenfrequcy of MD we apply simple scaling factor for the uniform spring constant:

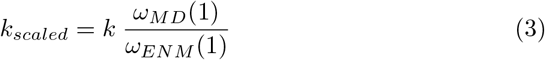

For BENM, alpha-carbon backbone spring values are multiplied by a scaling factor *ϵ*.

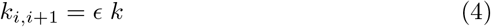

where *k*_*i,i*+1_ modified backbone spring constant between residues i^th^ and i+1^th^ on the same chain (See original work on BENM and allostery [26]).

### Cross-correlation of Motion

The cross-correlation for a mode *n, C*(*n*), is estimated between an ENM node pair as a normalised dot product between their normal mode eigenvectors.

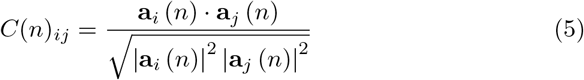

*C* value of 1 implies perfectly correlated motion, -1 perfectly anti-correlated motion and 0 implies totally non-correlated motion.

### Comparing ENMs with all-atom MD simulation

To find most suitable ENM distance cutoff we employed Chi-Square test for the first 25 non-trivial modes (n = 25) to compare ENM results with MD eigenfrequencies.

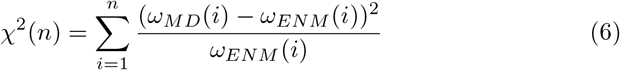

### Normal Mode Fluctuation Free Energy

Using statistical mechanics it is possible to calculate an estimate to the fluctuation free energy of a system using the frequency of vibrations such as the normal modes. For this method, the partition function for the quantum harmonic oscillator *Z*, for normal mode *n* is given as

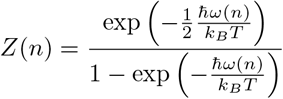

where *k*_*B*_ is the Boltzmann’s constant, *ћ* is the reduced Planck’s constant, *T* is temperature in Kelvin and *ω* is, already mentioned, angular frequency. The Gibbs free energy (for a given mode *n*) expressed in terms of partition function, with an approximation of little change in volume, can be written as

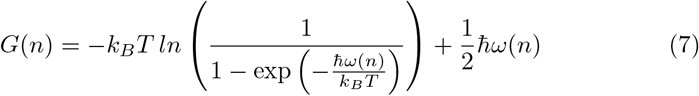

### Ligand Dissociation Constant

When the free energy change Δ*G* is known for a dissociation reaction, corresponding dissociation constant *K* can be estimated via

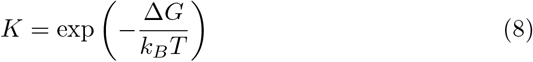

ENM predicted value of *K* was calculated by summing total number of mode *n* starting form the first non-trivial mode as follows

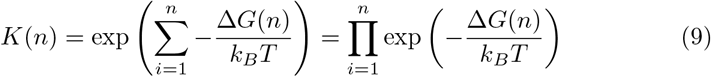

## Data accessibility

All MD, ENM, data processing and plotting scripts can be found https://github.com/tmcleish/article-optimising-enm as well as processed data used for the plots. Sufficient information and data are provided in and with the paper to allow others to replicate all study findings reported in this research article. Updated version of DDPT source code which includes backbone-enhancement can be accessed at https://github.com/tmcleish/ddpt.

## Acknowledgements

**ID** is grateful for computational support from the University of York high-performance computing service, the Viking Cluster. **TCBM** acknowledges support from the EPSRC (UK) through an Established Career Fellowship in the Physics of Life programme. **ID** and **TCBM** are both grateful to Prof Martin

J. Cann for suggesting GST as a protein to study and useful discussions.

