## Supplementary Information for "Optimising Elastic Network Models for Protein Dynamics and Allostery: Spatial and Modal Cut-offs and Backbone Stiffness"

### PDB structures

PDB files were processed by `pdb4amber` [1] and then trimmed from both chain ends to avoid floppy modes as indicated in table 1 of the article. `ATOM` records for ligands were change to `HETATM` so that DDPT could treat them as the part of a ligand, not a protein.
Atoms of bound molecules were manually removed from original PDB files (See Tab. 1) to make up the three protein forms: ligand-free (apo); singly-bound ligand to protomer A (holo1); each protomer, A and B, possess one ligand (holo2). PDB files for each protein used in this study can be found in the GitHub repository.

### DDPT

This section enumerates the routine used to generate and analyse proteins ENM via the Durham Dynamic Protein Toolbox (DDPT) [2]. As an example, the routine is carried out for a holo1 protein form (`01.pdb`). Apo and holo2 forms have file names of `00.pdb` and `02.pdb`, respectively. All programs from DDPT package are run with a command-line interface, also known as CLI.
First, the ENM interaction matrix is generated using the `GENENMM` program function in DDPT:

```
GENENMM -pdb 01.pdb -c 8 -het -b 50 -ca -mass -res -lig1 \
    -spcust spfile.ddpt -ccust cfile.ddpt
```

The crystallographic structure of a protein is assigned by `-pdb` flag. The cut-off distance is set by `-c`, by default 12 Å, and heteroatoms (`HETATM` in PDB notation) from the PDB file were included using `-het`, if required. `-b` adds backbone-enhancement factor to the ENM [3]. By default, all ENM nodes' masses (atoms from PDB structure are converted into ENM nodes) are set to the same fixed value, 1 amu, but actual atomic masses (`-mass`) were used in our ENM. The `-ca` flag creates an ENM model where amino acids' alpha-carbon atoms form nodes. In combination with `-ca`, the `-res` flag assigns the alpha-carbon nodes the whole residue mass. `-ccust` flag tells DDPT to read a passed file which specifies custom distance cutoff for residue types, while `-spcust` tells to read a file with custom bonds regardless the distance cutoff. The latter two flags are useful for ligand connectivity in an active because we might wish to connect only certain residues but not the others even if they are in a close proximity to the ligand. `GENENMM` program generates the interaction matrix for the ENM, written into `matrix.sdijf` file.

`-lig1` flag assigns a single node to each ligand (`HETATM`) based on mass-weighted ligand's atomic position (if `-mass` is absent than the resulting node coordinate is just an arithmetic average). Then, when `-res` is included, lignad's node is assigned its molecular mass which should be listed in `resmass.dat` file. This file must be present in the root directory where calculations take place (See DDPT manual).

Next, the `DIAGSTD` program diagonalises the interaction matrix using small-block diagonalisation and iterative schemes [2].

DIAGSTD -i matrix.sdijf

where `-i` flag specifies input Hessian matrix. The product of the diagonalisation is written into the `matrix.eigenfacs` file.

We calculate the inter-atom distance and cross-correlation of motion maps for the alpha-carbon ENM of the protein using `SPACING` and `CROSCOR` programs, respectively.

SPACING -pdb 01.pdb -ca

and

CROSCOR -i matrix.eigenfacs -s 7 -e 31

where `-s` and `-e` flags indicate the first and the last normal modes to include in calculation. Note, the first six normal mode frequencies, which correspond trivially to translational and rotational motion, are equal to zero.

Finally, fluctuation free energies for the set of the normal modes are calculated via the `FREQEN` program as follows

FREQEN -i matrix.eigenfacs -s 7 -e 106

The temperature ($T$) value used to calculate the partition function $Z$ and Gibbs free energy $G$ was 298 K, the default value for `FREQEN`.

### Fluctuation Free Energy Approximation

For convenience, DDPT estimates fluctuation free energy for each normal mode in dimensionless units $\frac{G}{k_B T}$ as follows

$$\frac{G}{k_B T} = -ln \left( \frac{1}{1 - \exp\left(-\frac{\hbar\omega}{k_B T}\right)} \right) + \frac{1}{2} \frac{\hbar\omega}{k_B T} \tag{1}$$

Normal mode frequencies of global modes in proteins are in the acoustic regime (less than 10 THz) [4, 5]. This fact is especially true for the slowest modes we investigated in this study. In this classical limit, $\dfrac{\hbar\omega}{k_B T} \ll 1$ at 298 K temperature. Therefore, `FREQEN` program in DDPT employs the following expression for fluctuation free energy calculation

$$\frac{G}{k_B T} \approx -ln \left( \frac{1}{1 - \exp\left(-\frac{\hbar\omega}{k_B T}\right)} \right) \tag{2}$$

In the differences ($\Delta G$) and difference of a difference ($\Delta\Delta G$) in fluctuation free energy (Eq. 3) significance of $\dfrac{1}{2} \dfrac{\hbar\omega}{k_B T}$ term is negligible.

### Fluctuation Energy Convergence

The allosteric free energy change is calculated using the fluctuation free energy change for three forms of the protein:

$$\begin{aligned} \Delta\Delta G &= \Delta G_2 - \Delta G_1 \\ &= (G_{holo2} - G_{holo1}) - (G_{holo1} - G_{apo}) \\ &= G_{holo2} - 2G_{holo1} + G_{apo} \end{aligned} \tag{3}$$

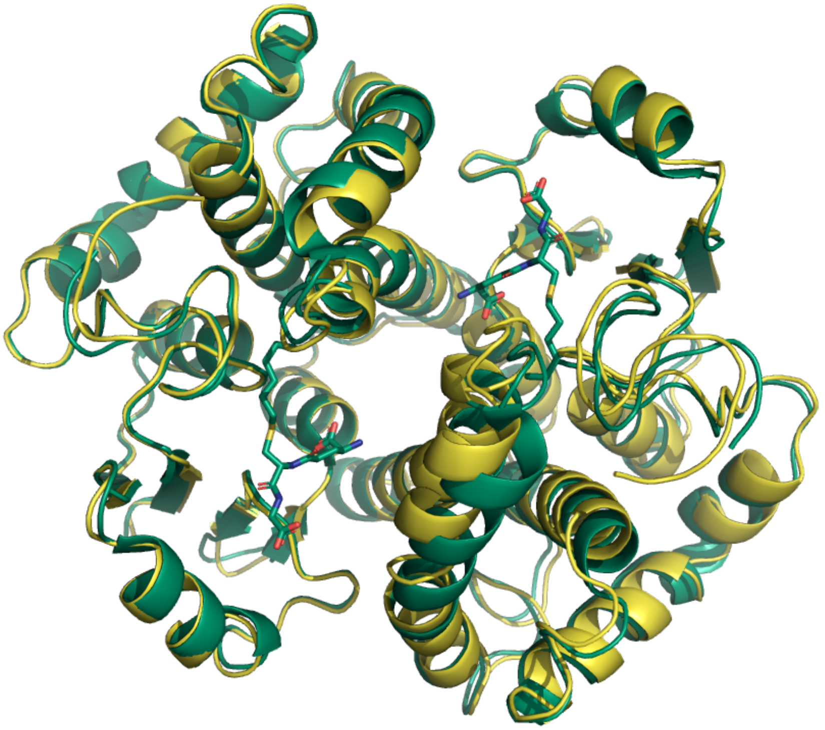

Figure 1: Alignment of *S. japonicum* GST apo form (PDB ID: 1GTA, *yellow*) and holo2 form with GTX in presented in *sticks* (PBD ID: 1M9A, *green*). Alignment performed in PyMOL using `cealign| algorithm yielded an RMSD of` 0.52 Å `for 216 residues.`

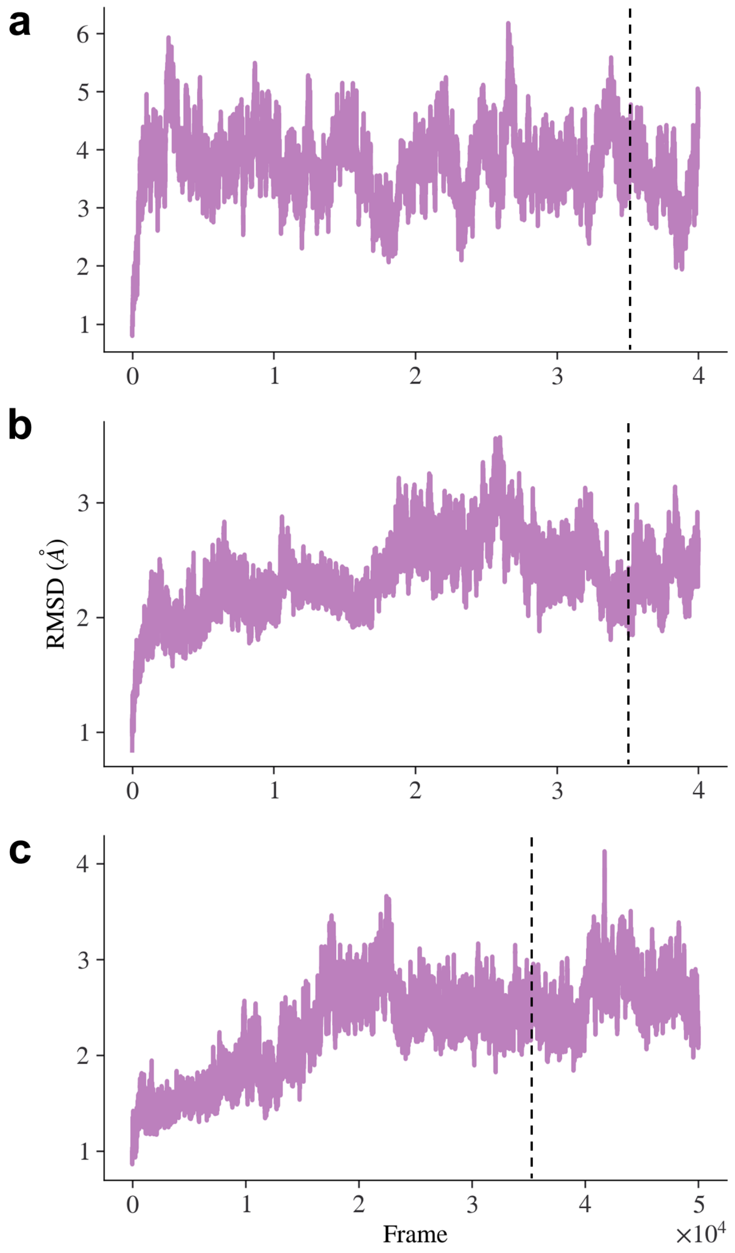


Figure 2: Equilibrating MD simulations. RMSD values from the beginning of MD simulations equilibration for CAP (**a**), GST (**b**) and $M^{pro}$ (**c**). The reference structure is X-ray derived structure from PDB. Terminal alpha-carbons are not included in the calculation. Each frames value is 0.01 ns. *Black* vertical dotted line indicates starting frame taken for PCA.

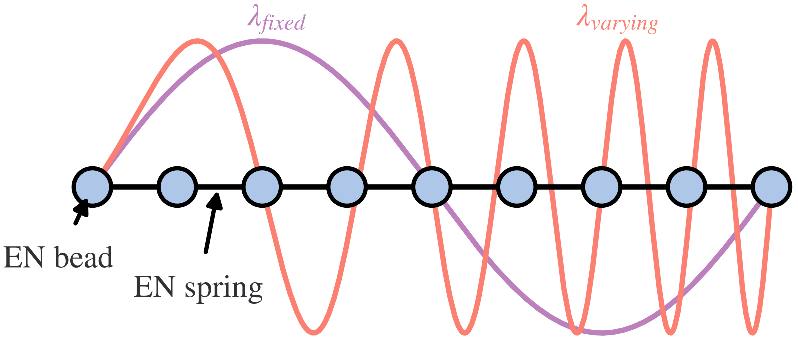


Figure 3: Normal modes in 1-dimensional elastic media. Two standing waves, purple and orange, with fixed and varying wavelength across the elastic network media, respectively. The higher density region (to the left) with the higher effective local elastic modulus supports a structure of the mode that is spatially resolved by the coarse-graining of the ENM at the level of nodes. However, the same mode contains a region of much finer structure (to the right in the diagram) which is not spatially resolved by the smallest length-scale offered by the ENM. Pale blue dots represent elastic network beads connected by Hookean springs.

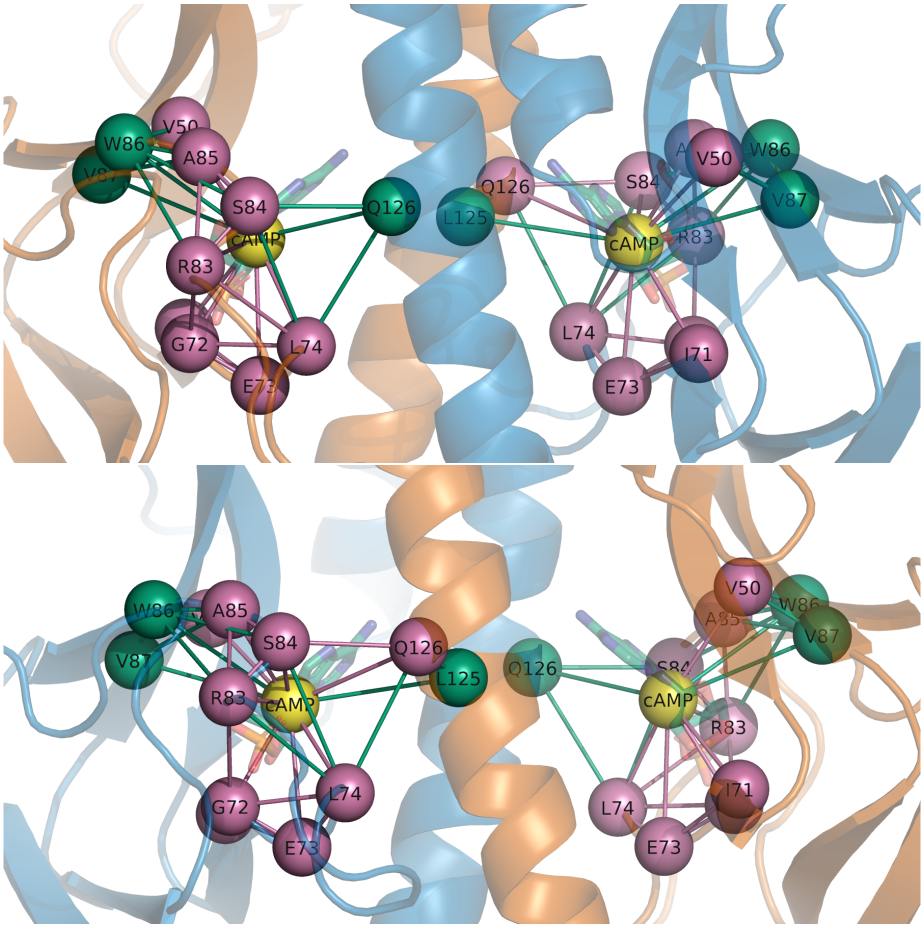


Figure 4: CAP's ENs comparison, (*top*) front and (*bottom*) back views, for $d_c = 7.5\,\text{Å}$ (*magenta*) and $d_c = 8\,\text{Å}$ (*green*). cAMP ligand node is coloured in *yellow*. CAP's cartoon protomer A and B are coloured in *orange* and **blue**, respectively. 7.5 Å EN has 8 EN springs attached to the ligand while 8 Å EN has 11.

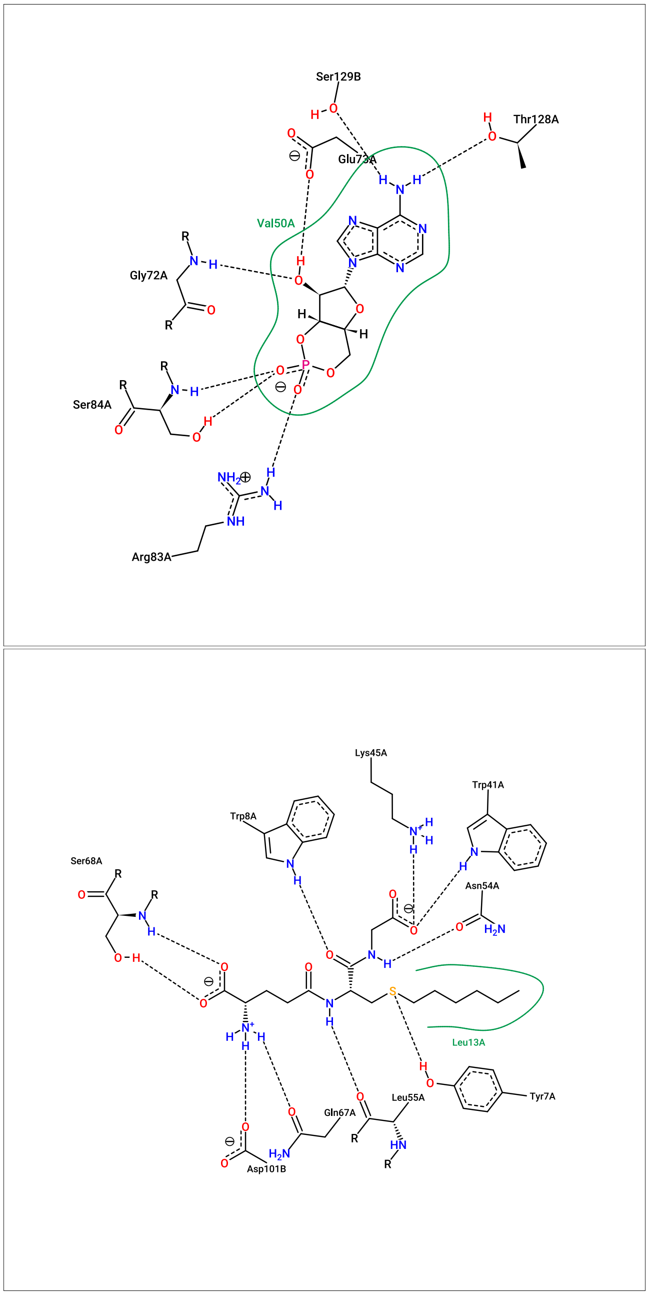


Figure 5: Protein-ligand connectivity based on X-ray structure for CAP (*top*) and GST(*bottom*). Both images were generated using online software tool PoseView [6]. Binding site information for SCoV2 M$^{pro}$ was taken from `https://www.rcsb.org/sequence/7BQY` (accessed May 12, 2022).

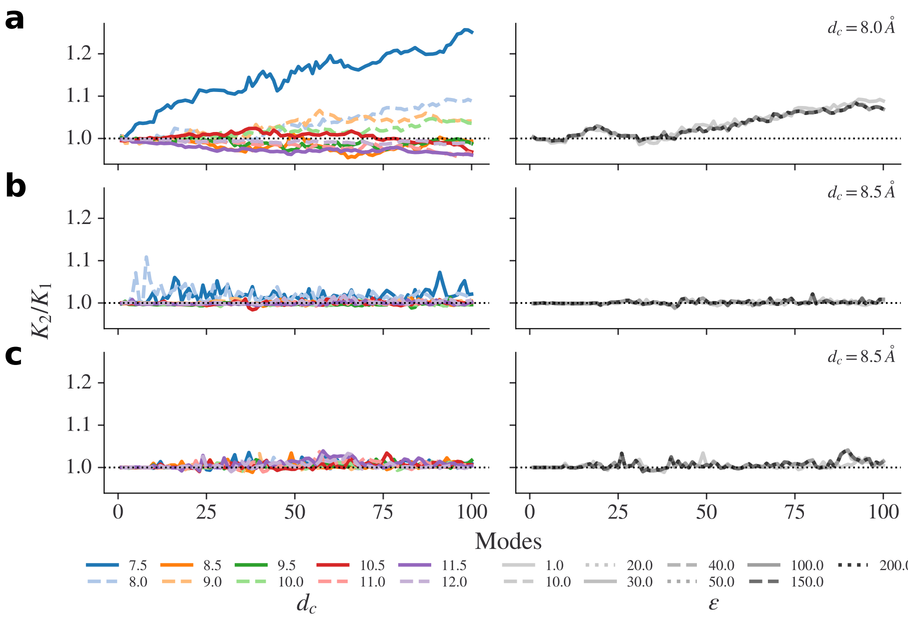


Figure 6: Cooperativity for ENM distance cutoff and BENM scan with a single-bead ligand and isotropic distance cutoff for it. Dotted line on the plots represent non-cooperative region. Each row presents data for a protein: **a** - CAP; **b** - GST; and **c** - $M^{pro}$. *Left* shows results for distance cutoff scan while *right* presents backbone-enhancement scan results.Each subplot represents the calculated cooperativity values against the total number of summed modes. Only first 100 non-trivial modes are shown. n each case, the ligand was modelled as a single mass-weighted elastic network bead with a total mass of the ligand.

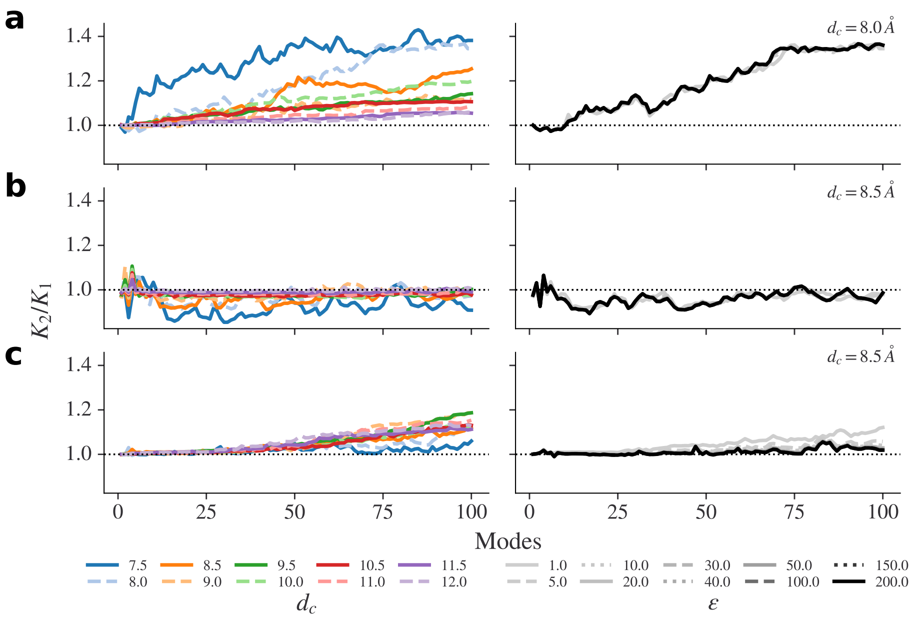


Figure 7: Cooperativity for ENM distance cutoff and BENM scan with ligand represented as a rigid body and custom protein-ligand bonds. Dotted line on the plots represent non-cooperative region. Each row presents data for a protein: **a** - CAP; **b** - GST; and **c** - $M^{pro}$. *Left* shows results for distance cutoff scan while *right* presents backbone-enhancement scan results. Each subplot represents the calculated cooperativity values against the total number of summed modes. Only first 100 non-trivial modes are shown. In each case, ligand heavy-atoms were treated as elastic network beads with according atomic mass and the ligand beads are all interconnected. The protein-ligand interaction was set as in the figure 9 of the main manuscript text: each selected protein residue connects to all ligand's beads.

**Algorithm 1** Pseudo-algorithm for the suggested in this study ENM selection based on all-atom MD simulation.

**1. Run** all-atom MD simulation for a given structure and perform PCA on the data.
**2. Scan** different ENMs with increasing distance cutoff $r_c$ values from 5 to 15 Å and discard ENMs with floppy modes.
$d_c = 5.0$ Å and $\Delta d_c = 0.5$ Å
**while** $d_c < 15.0$ Å **do**
    Calculate ENM with $d_c$.
    **if** $\omega(1) \approx 0$ **then**
        Discard current ENM.
    **end if**
    $d_c = d_c + \Delta d_c$.
**end while**
**3. Perform** $\chi^2$ test on ENMs and all-atom MD eigenvalues for the first 25 non-trivial modes and choose the best fit ENM.
